# Strong influence of vertebrate host phylogeny on gut archaeal diversity

**DOI:** 10.1101/2020.11.10.376293

**Authors:** Nicholas D. Youngblut, Georg H. Reischer, Silke Dauser, Chris Walzer, Gabrielle Stalder, Andreas H. Farnleitner, Ruth E. Ley

## Abstract

Commonly used 16S rRNA gene primers miss much of the archaeal diversity present in the vertebrate gut, leaving open the question of which archaea are host associated, the specificities of such associations, and the major factors influencing archaeal diversity. We applied 16S rRNA amplicon sequencing with Archaea-targeting primers to a dataset of 311 fecal/gut samples spanning 5 taxonomic classes (Mammalia, Aves, Reptilia, Amphibia, and Actinopterygii) and obtained from mainly wild individuals (76% were wild). We obtained sufficient archaeal sequence data from 185 samples comprising 110 species that span all 5 classes. We provide evidence for novel Archaea-host associations, including Bathyarchaeia and Methanothermobacter — the latter of which was prevalent among Aves and enriched in higher body temperatures. Host phylogeny more strongly explained archaeal diversity than diet, while specific taxa were associated with each factor. Co-phylogeny was significant and strongest for mammalian herbivores. Methanobacteria was the only class predicted to be present in the last command ancestors of mammals and all host species. Archaea-Bacteria interactions seem to have a limited effect on archaeal diversity. These findings substantially expand on the paradigm of Archaea-vertebrate associations and the factors that explain those associations.

**Significance:** Archaea play key roles in the vertebrate gut such as promoting bacterial fermentation via consumption of waste products. Moreover, gut-inhabiting methanogenic Archaea in livestock are a substantial source of greenhouse gas production. Still, much is not known of the archaeal diversity in most vertebrates, especially since 16S rRNA sequence surveys often miss much of the archaeal diversity that is present. By applying Archaea-targeted gut microbiome sequencing to a large collection of diverse vertebrates, we reveal new Archaea-host associations such as a high prevalence of Methanothermobacter in birds. We also show that host evolutionary history explains archaeal diversity better than diet, and certain genera in one particular class of Archaea (Methanobacteria) were likely pervasive in the ancestral vertebrate gut.

## Introduction

Next generation sequencing has greatly expanded our view of archaeal diversity, which now consists of nearly 40 major clades; 8 of which are currently known to be host-associated (1, 2). Many of these clades consist of methanogens, which utilize bacterial fermentation products (namely hydrogen and carbon dioxide) for obtaining energy and are generally the most abundant Archaea in the mammalian gut (3, 4). Halobacteria, Thaumarcheota, and Woesearchaeota comprise the major non-methanogenic host-associated archaeal clades and are generally not as prevalent or abundant among vertebrate gut microbiomes (2, 5).

Most data on archaeal diversity in the vertebrate gut derives from studies using standard “universal” 16S rRNA primers, which have recently been shown to grossly undersample archaeal diversity relative to using Archaea-targeting 16S rRNA gene primers (6–8). Therefore, much likely remains unknown of archaeal diversity and community assembly in the vertebrate gut. Setting primer issues aside, previous studies have identified host evolutionary history and diet to be main factors influencing the gut microbiome (9–13). While some studies have shown specific evidence that gut archaeal diversity is dictated by host relatedness (14–18), focus has generally been on humans and certain mammalian clades. Still, diet may also play a significant role, especially given that fiber can increase methanogen levels and ruminants generate substantial amounts of methane (3). Microbe-microbe interactions between Archaea and Bacteria may also have a strong influence on archaeal diversity, especially given the syntrophic interactions between methanogens and bacterial fermenters (19–21). Here, we characterize archaeal diversity in fecal/gut samples from 110 vertebrate species spanning 5 taxonomic classes. Using dietary and host phylogenetic relationships, as well as previously characterized bacterial diversity, we uncover robust relationships between Archaea, host phylogeny, and to some extent, host diet.

## Results

We utilized Archaea-targeting 16S rRNA primers that previously revealed vastly more gut archaeal community diversity in 5 great ape species relative to “universal” 16S primers (6). Our resulting gut microbiome 16S rRNA gene amplicon sequence dataset consisted of 185 samples from 110 species comprising 5 vertebrate classes (Figures 1 & S1). Most samples were derived from individual animals in the wild (76%), which is important given that captivity can alter the vertebrate gut microbiome (22, 23). Not all animal samples yielded adequate sequence data in order to be included in the final dataset (60% success; 185 of 311 samples), but failure was not correlated with host taxonomy, diet, or other characteristics (Figures S2 & S3).

**Figure 1.**
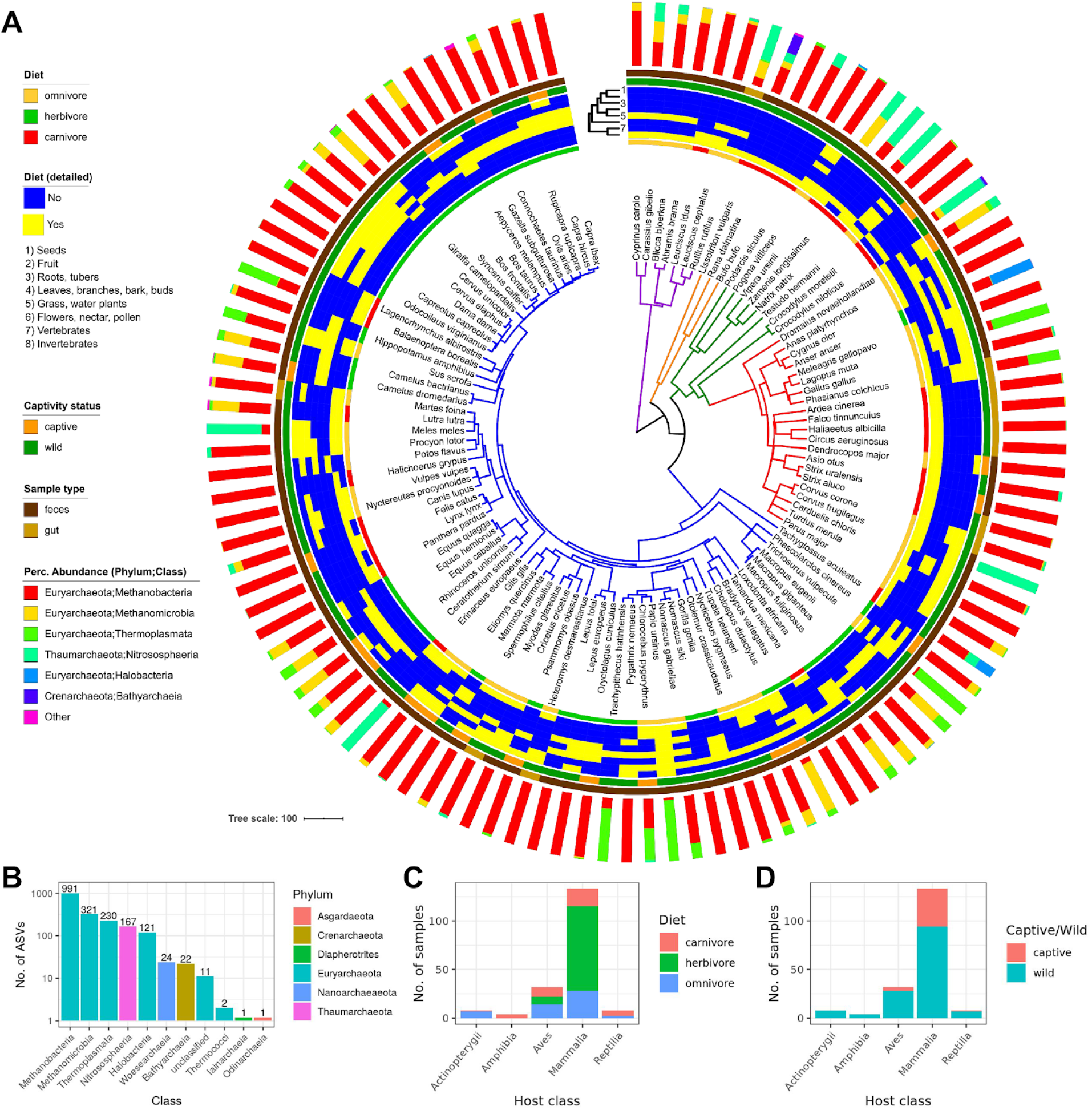
Substantial prevalence and diversity of Archaea among vertebrates. A) A dated phylogeny of all host species (*n* = 110) obtained from http://timetree.org, with branches colored by host class (purple = Actinopterygii; orange = Amphibia; green = Reptilia; red = Aves; blue = Mammalia). For inner to outer, the data mapped onto the phylogeny are: host diet (general), detailed diet composition (the dendrogram depicts Jaccard similarity), wild/captive status, sample type, and mean percent abundances of archaeal taxonomic classes among all individuals of the species. B) The number of ASVs belonging to each class. C) & D) The number of samples grouped by host class and C) diet or D) captive/wild status.

We found per-host archaeal diversity to be rather low, with only approximately 250 sequences saturating diversity estimates, regardless of host class or diet (Figure S4). Still, the taxonomic composition of the entire dataset was rather diverse for Archaea, comprising 6 phyla and 10 classes (Figure 1). The dataset consisted of 1891 amplicon sequence variants (ASVs), with dramatic phylum- and class-level compositional variation among host species but relatively low variation within species (Figure S5; Table S3). Methanobacteria (Euryarchaeota phylum) dominated in the majority of hosts. Thermoplasmata (Euryarchaeota phylum) dominated in multiple non-human primates, while two mammalian and one avian species were nearly completely comprised of Nitrososphaeria (Thaumarchaeaota phylum): the European badger (*Meles meles*), the Western European Hedgehog (*Erinaceus europaeus*), and the Rook (*Corvus frugilegus*). Halobacteria (Euryarchaoeota phylum) dominated the Goose (*Anser anser*) microbiome, which were all sampled from salt marshes. The class was also present in some distantly related animals (*e.g.*, the Nile Crocodile (*Crocodylus niloticus*) and the Short Beaked Echidna (*Tachyglossus aculeatus*)) (Tables S1 & S3).

Of the 10 observed archaeal classes, 4 are not known to include host-associated taxa (2): Bathyarchaeia, Iainarchaeia, Odinarchaeia, and Thermococci (Figure 1). The most prevalent and abundant was Bathyarchaeia, which comprised 9 ASVs present in 6 species from 4 vertebrate classes. It was rather abundant in the Nile Crocodile (3.3%) and the 2 Smooth Newt samples (17.9 and 42.2%) (Table S4). The other 3 classes comprised a total of 4 ASVs and were observed very sparsely and at low abundance, suggesting transience or persistence at very low abundances.

Only 40% of ASVs had a ≥97% sequence identity match to any cultured representative (Figure S6A). Of the 10 archaeal taxonomic classes, 5 had no match at ≥85% sequence identity: Odinarchaeia, Bathyarchaeia, Iainarchaeia, Woesarchaeia, and Thermococci. Taxonomic novelty to cultured representatives differed substantially among the other 5 classes but was still rather low (Figure S6B), even for relatively well-studied clades (*e.g.*, Methanobacteria). These findings suggest that our dataset consists of a great deal of uncultured taxonomic diversity.

Of 140 samples that overlap between our Archaea-targeted 16S dataset (“16S-arc”) and that from our previous work with standard “universal” 16S primers (“16S-uni”), 1390 versus only 169 archaeal ASVs were observed in each respective dataset (Figure S7). Representation of major clades was also much higher for the 16S-arc dataset. For example, Methanobacteria was observed in all host species via the 16S-arc primers, while prevalence dropped substantially for 16S-uni primers (*e.g.*, only 9% for Aves).

We used multiple regression on matrices (MRM) to assess the factors that explain archaeal diversity. Notably, we employed a permutation procedure to assess the sensitivity of our results to archaeal compositional variation among hosts of the same species (see Methods). Geographic distance, habitat, and technical components (*e.g.*, feces versus gut contents) did not significantly explain beta diversity, regardless of the diversity metric (Figure 2A). Host phylogeny significantly explained diversity as measured by unweighted UniFrac, Bray Curtis, and Jaccard (P < 0.05); however, significance was not quite reached for weighted UniFrac. The percent variation explained was dependent on the beta diversity measure and varied from ~28% for Jaccard to ~12% for unweighted UniFrac. In contrast to host phylogeny, composition of dietary components (diet) was only significant for Bray-Curtis, with ~12% of variance explained. Mapping the major factors onto ordinations qualitatively supported our results (Figure S8). Applying the same MRM analysis to just non-mammalian species did not generate any significant associations between host phylogeny or diet (Figure S9), likely due to the low sample sizes (*n* = 39). However, host phylogeny explained as much variance as when including all species, while variance explained by diet was relatively small. Altogether, these findings suggest that host evolutionary history mediates vertebrate gut archaeal diversity more than diet.

**Figure 2.**
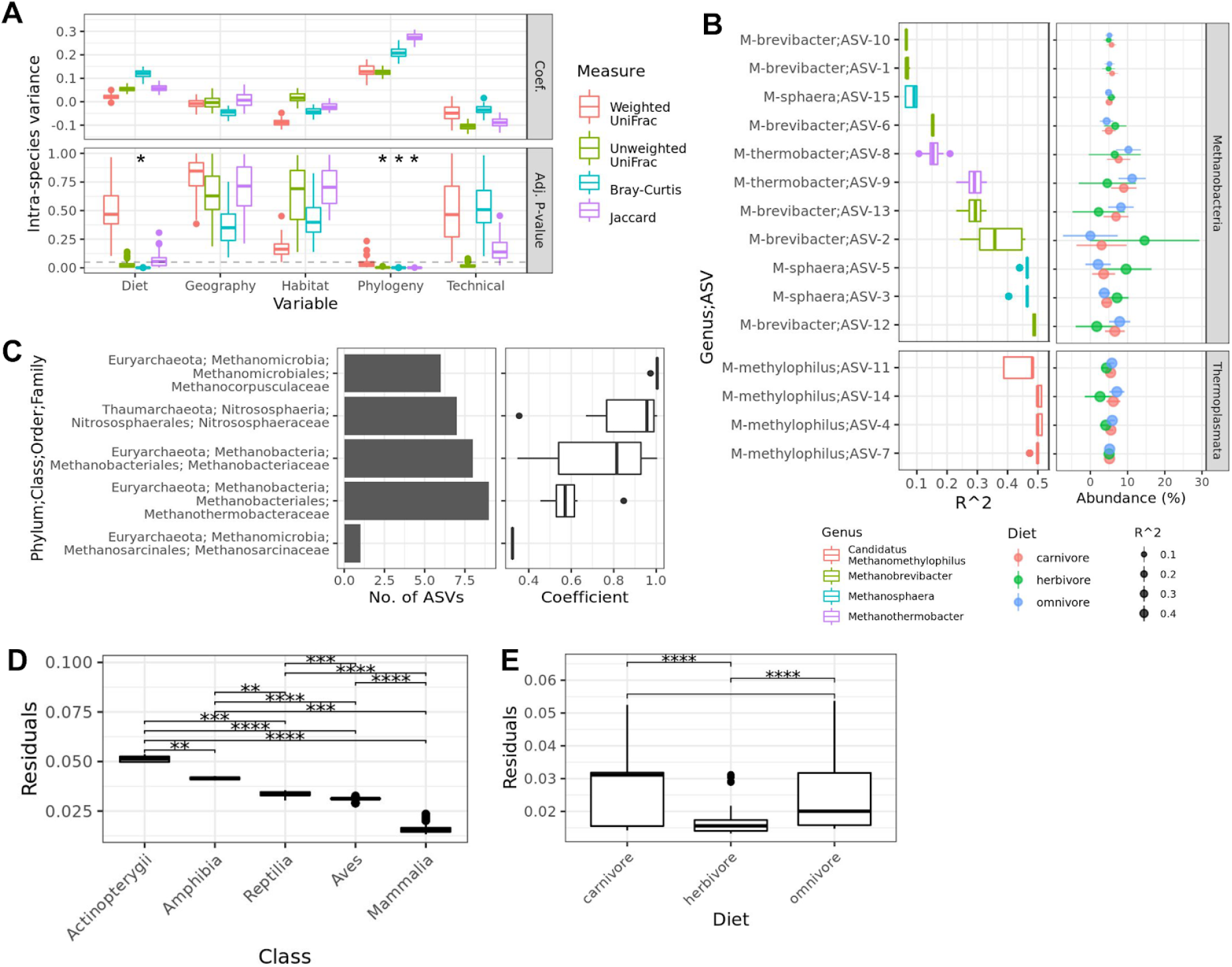
Host phylogeny and diet significantly explain different aspects of archaeal diversity. A) The distribution of partial regression coefficients (“Coef.”) and P-values (“Adj. P-value”) across 100 dataset permutations used for multiple regression on matrix (MRM) tests. For each permutation, one individual per host species was randomly sampled. MRM tests assessed the beta diversity variance explained by host diet, geography, habitat, phylogeny, and “technical” parameters (see Methods). Asterisks denote significance (adj. P < 0.05 for >95% of dataset subsets; see Methods). B) ASVs in which abundances are significantly correlated with diet (adj. P < 0.05) while controlling for host phylogeny via the randomization of residuals in a permutation procedure (RRPP). The left plot shows the distribution of coefficient values across all 100 permutations of the host tree, while the right plot shows RRPP model predictions of ASV abundances, depending on diet (points = mean; line ranges = 95% CI). C) The left plot shows the number of ASVs with a significant global phylogenetic signal (Pagel’s λ, adj. P < 0.05), while the right plot shows the distribution of coefficient values for those ASVs. D) & E) The distribution of PACo residuals across samples (averaged across all 100 dataset permutations) and grouped by D) host class or E) diet. Brackets indicate significant pairwise differences (Wilcox, ** < 0.01, *** < 0.001, **** < 0.0001). Box centerlines, edges, whiskers, and points signify the median, interquartile range (IQR), 1.5 × IQR, and >1.5 × IQR, respectively.

We also assessed alpha diversity via MRM in order to provide a consistent comparison to our beta diversity assessment (Figure S10). No factors significantly explained alpha diversity calculated via either the Shannon Index or Faith’s PD.

Although diet did not strongly explain total archaeal diversity, it may substantially explain the distribution of particular archaeal taxa. We used two methods to resolve the effects of diet on the archaeal microbiome while controlling for host evolutionary history: phylogenetic generalized least squares (PGLS) and randomization of residuals in a permutation procedure (RRPP) (24, 25). RRPP and PLGS identified the same 10 ASVs as being significantly associated with diet, while RRPP identified 5 more, likely due to increased sensitivity (adj. P < 0.05; Figures 2B & S11). All 15 ASVs belonged to the Euryarchaeota phylum and comprised 4 genera: Methanobrevibacter, Methanosphaera, Methanothermobacter, and candidatus Methanomethylophilus. RRPP model predictions of ASV abundances showed that Methanobacteria ASVs differed in their responses to diet, with 5 being most abundant in herbivores, while the other 6 were more abundant in omnivores/carnivores (Figure 2B). Notably, diet enrichment differed even among ASVs belonging to the same genus. In contrast to the Methanobacteria ASVs, all 4 Methanomethylophilus ASVs were predicted as more abundant in omnivores/carnivores. These findings suggest that diet influences the abundances of particular ASVs, and even closely related ASVs can have contrasting associations to diet. All significant ASVs were methanogens, which may be due to the species studied (*e.g.*, a mammalian bias) or possibly because certain methanogens respond more readily to diet.

When applied to alpha or beta diversity, neither PGLS nor RRPP identified any significant associations with diet (adj. P > 0.05). These findings correspond with our MRM analyses by indicating that diet is not a strong modulator of total archaeal diversity.

We also assessed whether particular archaeal taxa are explained by host evolutionary history and identified 37 ASVs to have abundances correlated with host phylogenetic relatedness (Pagel’s λ, adj. P < 0.05). These ASVs spanned three phyla: Euryarchaota, Thaumarchaeota, and Crenarchaeota (Figure 2C). The clade with the highest number of significant ASVs (*n* = 15) was Methanobacteriaceae, followed by Nitrososphaeraceae (*n* = 12), and Methanocorpusculaceae (*n* = 5). No such phylogenetic signal was observed when assessing alpha diversity rather than ASV abundances (adj. P > 0.05), which corresponds with our MRM results. We also tested for local instead of global correlations between ASV abundances and host phylogeny via the local indicator of phylogenetic association (LIPA). 25 ASVs showed significant associations with certain host clades (Figure S12A). For instance, 3 Nitrososphaeraceae ASVs were associated with 2 snake species (*Zamenis longissimus* and *Natrix natrix*), 3 Methanobrevibacter ASVs were associated with 2 species of kangaroo (*Macropus giganeus* and *Macropus fuliginosus*), and a Methanocorpusculum ASV was associated with both camel species (*Camelus dromedarius* and *Camelus bactrianus*). The 2 major exceptions to this trend were the Methanothermobacter ASVs, which associated with many species of Aves, while the Methanobrevibacter and Methanosphaera ASVs associated with many Artiodactyla species. Summarizing the number Archaea-host clade associations revealed clear partitioning of archaeal taxa by host clade, except for Methanobrevibacter, for which at least one ASV was associated with each of the host orders (*n* = 23 orders; Figure S12B).

To test for corresponding phylogenetic associations on both the host phylogeny and the archaeal 16S rRNA phylogeny, we employed two measures of co-phylogeny: Procrustes Application to Cophylogenetic Analysis (PACo) and ParaFit (26, 27). Both PACo and ParaFit tests were significant (P < 0.01). We found significant differences in the distribution of PACo Procrustes residuals among the vertebrate classes and diets (Kruskal-Wallis < 0.01; pairwise Wilcox < 0.01 for all), indicating stronger signals of cophylogeny in the order of (Mammalia > Aves > Reptilia > Amphibia > Actinopterygii) for taxonomy and (herbivores > omnivores/carnivores) for diet (Figure 2D & 2E).

We utilized ancestral state reconstruction (ASR) to investigate which archaeal clades were likely present in the ancestral vertebrate gut. Predictions of class-level abundances were accurate for extant hosts (adj. R^2^ = 0.86, P < 2e-16; Figure S14), and all 95% CIs were constrained enough to be informative (26% ± 29 s.d.). The model revealed that Methanobacteria was uniquely pervasive across ancestral nodes, while other classes were sparsely distributed among extant taxa and across a few, more recent ancestral nodes (Figures 3 & S15). Importantly, the model predicted that Methanobacteria was the only class to be present in the last common ancestor (LCA) of all mammals and the LCA of all 5 host taxonomic classes (Figures 3F & 3G). We generated a similar ASR model for all 4 Methanobacteria genera. Our model was more accurate at predicting extant traits than our class-level model (adj. R^2^ = 0.93, P < 2e-16; Figure S14) and 95% CIs were informative (28% ± 24 s.d.). The model predicted 3 of the 4 genera to be present in the LCA of all mammals and the LCA of all host species (Figure 3F & 3G). Methanobrevibacter and Methanothermobacter were predicted to have similar abundances for both LCAs (~30-35%), while Methanosphaera was much lower (~5%). The model predicted Methanobrevibacter to be most highly abundant in the Artiodactyla and generally abundant across most Mammalia clades (Figure S16), while Methanothermobacter was predicted to be most highly abundant and prevalent across the Aves and also mammalian clades in which Methanobrevibacter was less abundant (*e.g.*, Carnivora and Rodentia). Methanosphaera was prevalent across most animal clades, but generally at low abundance.

**Figure 3.**
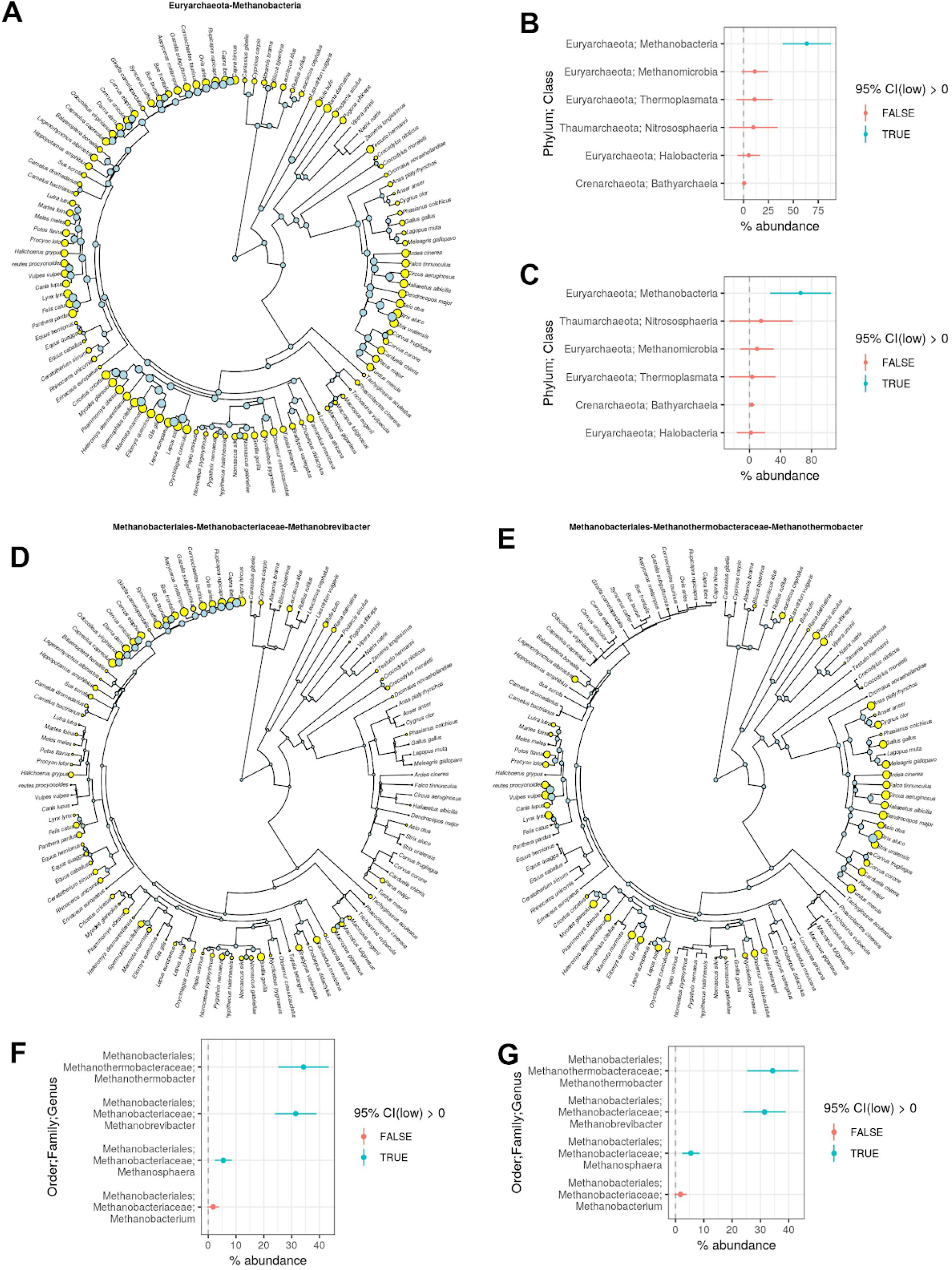
Ancestral state reconstruction evidences Methanobacteria association with ancestral vertebrate gut. A) Predicted abundances of Methanobacteria for each extant host species (yellow circles) and ancestral host species (blue circles). Circle size denotes relative abundance (min = 1%, max = 100%). The phylogeny is the same as shown in Figure 1. B) & C) Estimated Methanobacteria abundance (points) with 95% CIs (line ranges) for the LCA of B) all mammalian species and C) all 5 taxonomic classes. D) & E) The same as A), but predicted abundances of the Methanobrevibacter and Methanothermobacter genera, respectively. The plots in F) and G) are the same as B) and C), respectively, but show predictions of abundances for all 4 genera in the Methanobacteria class.

Methanothermobacter is not known to be host-associated (2). Still, a total of 39 Methanothermobacter ASVs were observed across 78 samples (18 ± 30 s.d. samples per ASV), which strongly suggests that its presence is not due to contamination. Moreover, the top BLASTn hit for 36 of the 39 ASVs was to a cultured Methanothermobacter strain (Figure S17, Table S5), which indicates that the taxonomic annotations are demonstrably correct. The high prevalence of Methanothermobacter among Aves lead us to the hypothesis that body temperature significantly affects the distribution Methanothermobacter (Figure S18), given that birds generally have higher body temperatures than mammals (28) and all existing Methanothermobacter strains are thermophiles (29). Moreover, Methanothermobacter is not abundant in Monotremata and Marsupialia species relative to the placental groups, which reflects the higher body temperature of placentals (Figure S18). We were able to assign published body temperature data to 73 mammalian and avian species (Figures S19A & S19B; Table S6). Genus-level abundances of Methanothermobacter significantly correlated with body temperature (RRPP, adj. P < 0.001), while Methanobrevibacter and Methanosphaera did not (Figures S19C & S19D). However, the association was only significant if not accounting for host phylogeny (RRPP, adj. P > 0.05), indicating that the association between Methanothermobacter and body temperature could not be decoupled from host evolutionary history. We also identified 7 Methanothermobacter ASVs to be correlated with body temperature (RRPP, adj. P < 0.05; Figure S19E), while no Methanobrevibacter or Methanosphaera ASVs were correlated. Again, the association was only significant if not accounting for host phylogeny. Regardless, we provide evidence congruent with the hypothesis that Methanothermobacter abundance is modulated by host body temperatures. We note that among the host species in which methane emission data exists (14, 30), avian species with high abundances of Methanothermobacter have emission rates on the higher end of mammal emission rates (Figure S20), suggesting that Methanothermobacter is indeed a persistent inhabitant in the gut of some avian species.

Besides host-specific factors modulating diversity, microbe-microbe interactions may also play a significant role. A co-occurrence network of all archaeal ASVs revealed high assortativity by taxonomic group, regardless of the taxonomic level (Figures S21 & S22). The only significant negative co-occurrences were between a cohort dominated by Methanobrevibacter and one dominated by Methanothermobacter. These 2 cohorts differed substantially in their distributions across host clades, with the Methanobrevibacter-dominated cohort only highly prevalent among Artiodactyla, while the Methanothermobacter-dominated cohort was prevalent across a number of mammalian orders (*e.g.*, Carnivora and Rodentia) and almost all avian orders (Figure S23). We assessed whether taxonomic assortativity differs among host diets (Figure S24), and indeed we found assortativity to be lowest for omnivores and highest for carnivores, suggesting that the carnivore gut is composed of simpler and more taxonomically homogenous archaeal consortia relative to omnivores and herbivores.

We assessed Archaea-Bacteria interactions by comparing the overlapping 16S-arc and 16S-uni samples (*n* = 140). Archaeal and bacterial alpha and beta diversity were not correlated, regardless of measure (adj. P > 0.05; Figure 4A & 4B), which suggests that total archaeal diversity is not explained by bacterial diversity nor *vice versa*. Overall network taxonomic assortativity was low; however, assortativity of just Archaea was quite high (≥0.774 for all taxonomic levels; Figure 4E). As with the Archaea-only network, there were 2 cohorts dominated by different Archaea: the first by Methanobrevibacter and then second by Methanothermobacter (Figure 4C & 4D). These 2 cohorts differed dramatically in bacterial diversity. The Methanobrevibacter cohort comprised 13 bacterial families from 3 phyla, while the Methanothermobacter cohort included only 3 families from 2 phyla (Figure S25).

**Figure 4.**
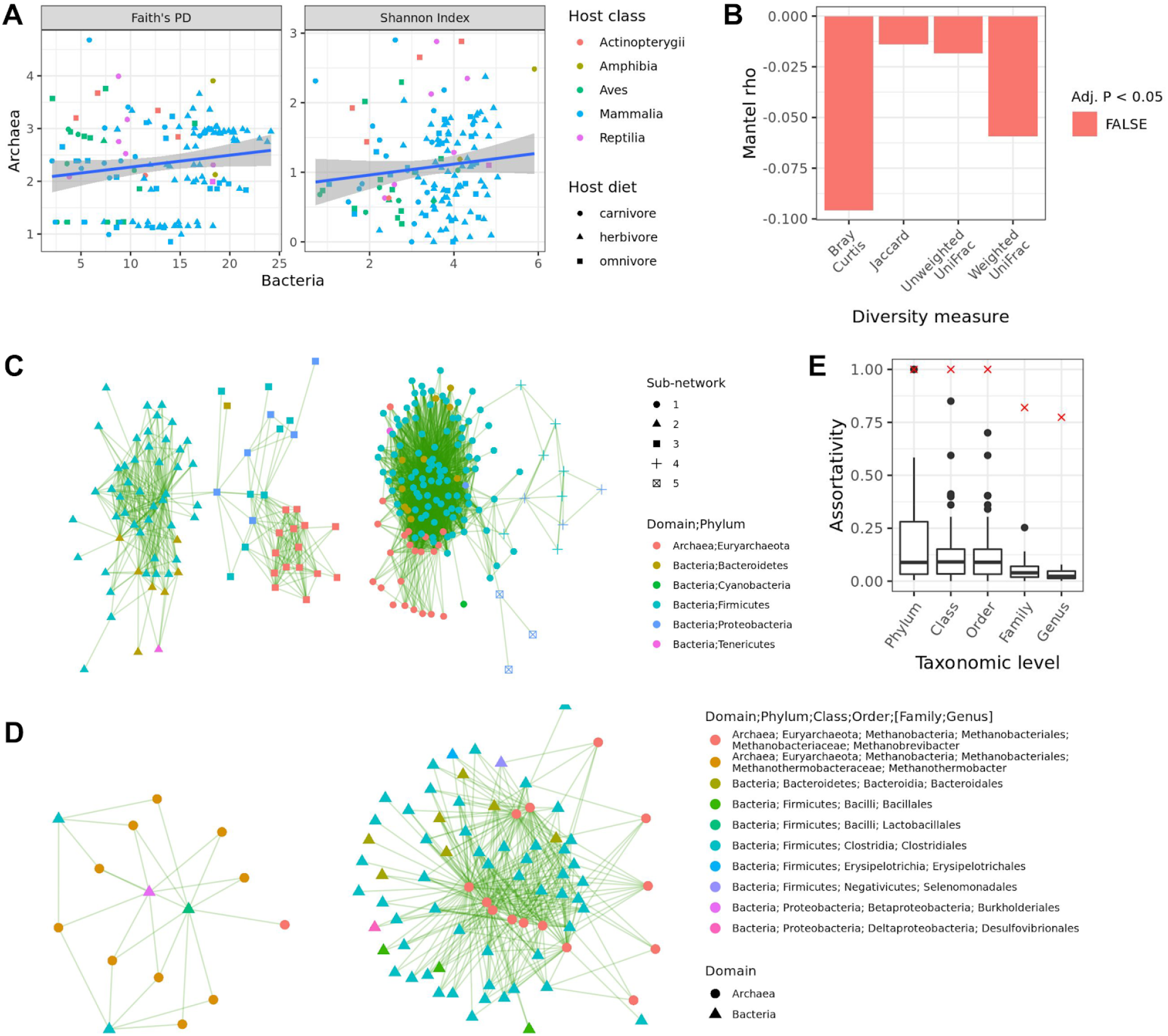
Limited associations between archaeal and bacterial diversity. A) Linear regressions of archaeal versus bacterial community diversity as measured via Faith’s PD and the Shannon Index. The grey areas denote 95% CIs. B) Mantel tests comparing archaeal and bacterial beta diversity (999 permutations per test). C) Archaea-Bacteria ASV co-occurrence network, with nodes colored at the phylum level. Green edges indicate significant positive co-occurrences. D) Just the sub-networks of the co-occurrence network shown in C) that contain archaeal ASVs. E) Taxonomic assortativity of just Archaea-Archaea edges (red “X”) versus just Bacteria-Bacteria edges (box plots), with Bacteria subsampled to the same number of ASVs as Archaea (100 subsample permutations). Box centerlines, edges, whiskers, and points signify the median, interquartile range (IQR), 1.5 × IQR, and >1.5 × IQR, respectively.

## Discussion

We show that the vertebrate gut harbors a great deal more archaeal diversity than previously observed (Figures 1 & S7) (13, 22). Although we were unsuccessful at obtaining sufficient archaeal amplicon sequence data from many samples (60% success rate), the broad host taxonomic diversity of our final dataset suggests that Archaea are widespread among vertebrata (Figures 1, S2, & S3). Moveover, the diversity of archaeal ASVs across samples and repeated intra-host-species observations of archaeal taxa indicate that the vertebrate gut microbiome collectively harbors a diverse archaeal assemblage.

Little is known of the gut microbiome for many of the host species in our dataset, especially regarding archaeal diversity. Only a minority of all known archaeal phyla and classes include any cultured representative (1), and accordingly we show that the majority of ASVs in our dataset lack cultured representatives (Figures S6 & S17), even ASVs belonging to more well-studied archaeal clades (*e.g.*, Methanobrevibacter). Interestingly, our dataset included repeated observations of archaeal clades not known to inhabit the vertebrate gut (2), suggesting previously unknown Archaea-vertebrate associations. For instance, the Bathyarchaeia (also known as candidatus Bathyarchaeota) is not known to include host-associated members (2), but this clade consisted of 9 ASVs and was substantially abundant in multiple individuals comprising various, distantly related species (Figure 1). Bathyarchaeia currently lacks any cultured representatives (1), but inference from metagenome-assembled genomes suggests that the clade contains methanogens and homoacetogens, and has been detected across a wide range of environments, especially aquatic biomes (31, 32). The relatively high abundance of Bathyarchaeia in the Smooth Newt and the Nile Crocodile, both of which have semi-aquatic lifestyles, may suggest adaptation to the gut from sediment-inhabiting ancestors. Alternatively, consumption of sediment may result in its transitory presence in the vertebrate gut, but one would then expect more Bathyarchaeia presence in other (semi-)aquatic vertebrates (*n* = 24) versus what we observed (Figure 1; Table S4).

Methanothermobacter is another clade not known to be host-associated, especially given the high optimal growth temperature (55 - 66°C) of existing cultures (29, 33). Regardless, we observed a diverse set of Methanothermobacter ASVs that were most prevalent and abundant among avian species (Figure S18). How is Methanothermobacter inhabiting the relatively cool gut environment, and why is this clade not considered host-associated? First, some Methanothermobacter isolates can grow at 37°C, but not optimally (33). Second, Methanothermobacter has been observed in a number of animals including salamanders (34), trout (35), chickens (36), buffalo (37) but very sparsely and at low abundances. Third, we found Methanothermobacter abundances to correlate with higher body temperatures of avian and some mammal species which have generally been less-studied, especially in regards to archaeal diversity (Figures S18 & S19).

We identified factors that significantly explain inter-species variation in archaeal community composition. Overall, habitat did not significantly explain inter-species beta diversity (Figure 2A); however, the high abundance of halophilic archaea in host species inhabiting high saline habitats (*e.g.*, geese) hints at habitat playing a role in certain cases (Figure 1). Both host phylogeny and diet correlated with beta diversity, but host phylogeny was more consistently significant among diversity metrics and showed stronger correlations (Figure 2A). These findings suggest that host evolutionary history has a stronger effect on total archaeal community diversity among host species relative to diet (both general dietary categories and specific components), which corresponds with existing evidence of an association between archaeal diversity and host phylogeny (12, 14). Still, these results contrast our previous Bacteria-biased survey of the same sample dataset, in which diet was an equal if not stronger explanatory variable relative to host phylogeny (13).

This difference between how diet associates with bacterial and archaeal diversity may be a result of 3 major factors. First, methanogens dominate most archaeal communities in our dataset (Figure 1), and these Archaea may not respond readily to diet since they only secondarily consume a relatively homogenous set of bacterial fermentation products (2, 4). Second, the nitrifying Archaea that dominate certain archaeal communities may be strongly influenced by the nitrogenous content of dietary components, which we did not directly measure. For instance, very high abundances of Thaumarchaeaota in certain vertebrates (*i.e.*, the European badger, the Western European Hedgehog, and the Rook) may be the result of consuming invertebrates such as earthworms, pill bugs, and termites; all of which all have been observed to harbor gut-inhabiting Thaumarchaeaota (38–40). Third, diet may only strongly influence a minority of archaeal ASVs, given that we only observed significant associations between ASV abundances and host diet for 15 ASVs (Figures 2B & S11). Interestingly, closely related Methanobacteria ASVs showed contrasting associations with diet, suggesting niche partitioning at fine taxonomic scales, possibly due to differing syntrophic associations with bacterial fermenters. Indeed, the pan-genome of *M. smithii* is highly variable in adhesin-like proteins thought to mediate cell-cell interactions with specific bacterial syntrophs (16, 41).

While previous work has indicated that host phylogeny influences archaeal abundance (12–14, 30, 42), the work has largely been focused on methanogen diversity (especially Methanobrevibacter) and was strongly biased towards mammals. In this work, we not only identified a signal of host-Archaea phylosymbiosis while accounting for host diet, habitat, and other factors (Figure 2A), but we also identified specific archaeal ASVs to be associated with host phylogeny globally and for particular host clades (Figures 2C & S12), and we observed a significant signal of cophylogeny (Figure 2D & 2E). While both the signal of phylosymbiosis and cophylogeny may be dominated by methanogens, we did identify specific non-methanogenic ASVs to be associated with host phylogeny (Figure S12), indicating that the influence of host phylogeny is not restricted to methanogens. These non-methanogenic ASVs were all Thaumarchaeota and showed host clade specificity to reptiles and bony fish (Figure S12), which emphasizes a need to study non-methanogenic Archaea outside of mammals and birds. Interestingly, all significant local phylogenetic signals for mammals were herbivorous species. These results concur with our assessments of co-phylogeny, which show the strongest signal for mammals and herbivores (Figures 2D & 2E). We previously observed a similar result when focusing on the bacterial community (13), suggesting that both host-Archaea and host-Bacteria co-phylogeny is strongest for herbivorous mammals. Many characteristics typical of mammals may be driving this pattern, such as complex gut morphology and placental birth (43, 44).

Based on methane emission data, Hackstein and van Alen hypothesized that methanogens were present in the gut of the Mammalia LCA and likely even present in the Vertebrata LCA, but certain lineages have permanently lost this “trait” (14, 42). Our ancestral state reconstruction of archaeal abundances did indeed support methanogen presence in the LCA of mammals and all 5 taxonomic classes of vertebrates (Figure 3), but importantly, we specifically identified Methanobacteria as the only archaeal class to show such evidence. Moreover, we evidenced 3 of the 4 Methanobacteria genera (*i.e.*, Methanobrevibacter, Methanothermobacter, and Methanosphaera) to be present in the LCA of mammals and all 5 classes (Figure 3). The prevalence of these 3 genera across all vertebrate classes suggests that methanogens are not completely lost from certain lineages but rather just differ in relative abundance (Figure 3). The relatively high predicted abundance of Methanothermobacter in the LCA of all host species may be biased due to its high abundance in most avian species and the limited number of ectothermic host species. Nevertheless, our findings suggest a long evolutionary association between vertebrates and certain methanogens.

We found little evidence of microbe-microbe interactions influencing archaeal diversity. Archaea generally assorted with sister taxa, regardless of diet (Figure S24), which may be the result of strong selective pressure counteracting competitive exclusion. Indeed, we observed niche partitioning by diet among closely related Methanobrevibacter ASVs (Figures 2B & S11). Moreover, closely related Archaea may be spatially partitioned along the GI tract as observed in human biopsies (7), or along other niche axes such as virus-Archaea interactions (45). Archaeal taxonomic assortativity was high relative to Bacteria (Figure 4E), indicating that community assembly processes differ between the domains, which corresponds with the lack of correlation between archaeal and bacterial diversity (Figures 4A & 4B). These findings indicate that methanogen-Bacteria syntrophic interactions do not strongly dictate archaeal diversity (2, 4, 46). Interestingly, we observed 2 major Archaea-Bacteria assemblages dominated by Methanobrevibacter and Methanothermobacter, respectively. These assemblages reflect the different distributions of each methanogen genus, which is likely influenced by body temperature (Figures S18 & S19). Given that members of both methanogen clades are hydrogenotrophs (29), possess adhesins for attachment to organic surfaces (41, 47), and share many homologs via horizontal gene transfer and shared common ancestry (37), body temperature may be a main niche axis partitioning these two clades.

In conclusion, our findings expand the paradigm of Archaea-host interactions, including novel host-Archaea associations for clades previously considered to not be host-associated (*e.g.*, Methanothermobacter). More work is needed to elucidate the role of host body temperature in modulating archaeal community assembly in the vertebrate gut, especially in regards to Methanothermobacter and Methanobrevibacter. Moreover, little is known of the potential host and microbial genetic factors underlying our observed signals of phylosymbiosis and cophylogeny. One promising approach is to combine genome-capture for obtaining genome assemblies of rare microbial taxa (48) with microbiome genome-wide association analyses (mGWAS) focused on archaeal strains (49).

## Materials and Methods

Sample collection was as described by Youngblut and colleagues (13). PCR amplicons for the V4 region of the 16S rRNA gene were generated with primers arch516F-arch915R (6, 50) and were sequenced with the Illumina MiSeq 2 × 250 v2 Kit. The host phylogeny was obtained from http://timetree.org (51). For all hypothesis testing (unless noted otherwise), we generated 100 permutation datasets in which one sample was randomly selected per host species, and a hypothesis test was considered robustly significant if >95% of the permutation datasets generated a significant result (P < 0.05 unless otherwise noted). See the supplemental Materials and Methods for all methodological details.

The raw sequence data are available from the European Nucleotide Archive under the study accession number PRJEB40672. All sample metadata used in this study is provided in Table S1. All code and the software versions used for analyses are available at https://github.com/leylabmpi/16S-arc_vertebrate_paper.

## Acknowledgements

This study was supported by the Max Planck Society and the Austrian Science Fund (FWF) research projects P23900 granted to Andreas H. Farnleitner and P22032 granted to Georg H. Reischer. Further support came from the Science Call 2015 “Resource und Lebensgrundlage Wasser” Project SC15-016 funded by the Niederösterreichische Forschungs-und Bildungsgesellschaft (NFB).

We thank Andre Teare for providing body temperature data from the Medical Animal Records Keeping System (MedArks). We thank the following collaborators for their huge efforts in sample and data collection: Mario Baldi, School of Veterinary Medicine, Universidad Nacional de Costa Rica; Wolfgang Vogl and Frank Radon, Konrad Lorenz Institute of Ethology and Biological Station Illmitz; Endre Sós and Viktor Molnár, Budapest Zoo; Ulrike Streicher, Conservation and Wildlife Management Consultant, Vietnam; Katharina Mahr, Konrad Lorenz Institute of Ethology, University of Veterinary Medicine Vienna and Flinders University Adelaide, South Australia; Peggy Rismiller, Pelican Lagoon Research Centre, Australia; Rob Deaville, Institute of Zoology, Zoological Society of London; Alex Lécu, Muséum National d’Histoire Naturelle and Paris Zoo; Danny Govender and Emily Lane, South African National Parks, Sanparks; Fritz Reimoser, Research Institute of Wildlife Ecology, University of Veterinary Medicine Vienna; Anna Kübber-Heiss and Team, Pathology, Research Institute of Wildlife Ecology, University of Veterinary Medicine Vienna; Nikolaus Eisank, Nationalpark Hohe Tauern, Kärnten; Attila Hettyey and Yoshan Moodley, Konrad Lorenz Institute of Ethology, University of Veterinary Medicine Vienna; Mansour El-Matbouli and Oskar Schachner, Clinical Unit of Fish Medicine, University of Veterinary Medicine; Barbara Richter, Institute of Pathology and Forensic Veterinary Medicine, University of Veterinary Medicine Vienna; Hanna Vielgrader and Zoovet Team, Schönbrunn Zoo; Reinhard Pichler, Herberstein Zoo. We explicitly thank the Freek Venter of South African National Parks and the National Zoological Gardens of South Africa for granting access to their Parks for sample collection.

## Author Contributions

G.H.R., R.E.L., and A.H.F. created the study concept. G.H.R., N.S., C.W., and G.S. performed the sample collection and metadata compilation. G.H.R., N.S., and S.D. performed the laboratory work. N.D.Y. performed the data analysis. N.D.Y. and R.E.L. wrote the manuscript.

