## Supplemental Materials for "Strong influence of vertebrate host phylogeny on gut archaeal diversity"

### Supplemental Materials and Methods

#### *Sample collection*

Sample collection was as described by Youngblut and colleagues (Youngblut et al. 2019). Samples used in this study were collected between February 2009 and March 2014. Only fresh samples with confirmed origin from a known host species were collected. Table S1 lists all dates, locations, and other relevant metadata associated with each sample. All fecal samples were collected in sterile sampling vials, transported to a laboratory and frozen within 8 hours. DNA was extracted with the PowerSoil DNA Isolation Kit (MoBio Laboratories, Carlsbad, USA).

#### *16S rRNA gene sequencing and data processing*

PCR amplicons for the V4 region of the 16S rRNA gene were generated with primers arch516F-arch915R (Takai and Horikoshi 2000; Raymann et al. 2017) and were sequenced with the Illumina MiSeq 2 × 250 v2 Kit at the Max Planck Institute for Developmental Biology. DADA2 (Callahan et al. 2016) was used to generate amplicon sequence variants (ASVs). Taxonomy was assigned to ASVs with the QIIME2 q2-feature-classifier (Bokulich et al. 2018) using the SILVA database (v119) (Pruesse et al. 2007). All ASVs not classified as Archaea were removed. Rarefaction analysis using alpha diversity quantified via the Vegan R package (Shannon Index; (Oksanen et al. 2012)) or the iNEXT R package (Hill numbers: order = 1; (Hsieh, Ma, and Chao 2016)) revealed that archaeal diversity saturated at a sampling depth of approximately 250 (Figure S4). Therefore, the dataset was rarefied to this depth, with all samples lacking this depth filtered out. Due to the low prevalence of ASVs across host species (1.8% ± 23 s.d.), we did not employ the standard compositional data analysis transformation of centered log ratio (CLR), given the large number of zero values in the dataset that would need to be imputed as non-zero values prior to the transformation. We found such imputation by either using a pseudo count of 1 or imputing via the Bayesian-multiplicative replacement method implemented in the zCompositions (Palarea-Albaladejo and Martín-Fernández 2015) R package generated unrealistic distributions. QIIME2 was used to calculate alpha and beta diversity. To limit saturation of star-phylogeny beta diversity measures (*i.e.,* no overlap of any ASVs across samples leading to maximum diversity values), we first aggregated ASV counts at the genus-level. A phylogeny was inferred for all ASV sequences with fasttree (Price, Dehal, and Arkin 2010) based on a multiple sequence alignment generated by mafft (Katoh and Standley 2013). All samples lacking relevant metadata used in the study were filtered from the dataset. In cases where an individual host was sampled multiple times, we randomly selected one sample.

Samples from the 16S rRNA amplicon dataset of Youngblut and colleagues were previously sequenced and process in the same manner as done for the arch516F-arch915R amplicon dataset, with the exception that the primers 515F-806R were used and samples were rarefied to a depth of 5000 (Youngblut et al. 2019). To compare ASVs classified as Archaea in each dataset, we filtered out all non-archaeal ASVs. For our analyses of Bacteria-Archaea interactions, we removed all archaeal ASVs from the 515F-806R dataset. Alpha and beta diversity were calculated as stated above on genus-level abundances.

*Host phylogeny*

Only 21% of animals in our dataset have existing genome assemblies of any quality in which to infer a genome-based phylogeny from. Instead, we used a dated host phylogeny for all species from [http://timetree.org](http://timetree.org) (Kumar et al. 2017). We created a phylogeny for all samples by grafting sample-level tips into each species node with a negligible branch length (Figure S5).

*Intra-species sensitivity analysis*

The dataset consisted of a differing number of samples per host species and no intra-species phylogenetic relatedness data. Instead of just randomly subsampling one sample per species or using branches of zero length for phylogeny-based hypothesis testing, we instead employed a sensitivity analysis to assess robustness to intra-species variability. The sensitivity analysis was performed as described in Youngblut and colleagues (Youngblut et al. 2019). Briefly, for each hypothesis test, we generated 100 permutation datasets in which one sample was randomly selected per species. A hypothesis test was considered robustly significant if >95% of the permutation datasets generated a significant result ($P < 0.05$ unless otherwise noted).

*Data analysis*

We used BLASTn (Camacho et al. 2009) to assess similarity of ASVs to cultured representatives in the SILVA All Species Living Tree database (Quast et al. 2013), with an E-value cutoff of <1e-5. All BLAST hits with an alignment length <95% of the query sequence length were filtered out.

Multiple regression on matrices (MRM) was performed with the Ecodist R package (Goslee and Urban 2007). We used rank-based correlations and 999 permutations to ascertain test significance. Regression variables that were not inherently distance matrices were converted via various means. Gower distance was used to convert detailed diet data, detailed habitat data, and “technical” data (*i.e.,* captive/wild animal and feces/gut-contents sample type) to distance matrices. Geographic distance was calculated as Great Circle distance based on sample latitude and longitude. Alpha diversity was converted to a Euclidean distance matrix. Principal coordinate analysis (PCoA) ordinations were generated for each beta diversity measure via the Vegan R package (Oksanen et al. 2012).

Pagel’s $\lambda$ and the local indicator of phylogenetic association (LIPA) were calculated via the Phylosignal R package (Keck et al. 2016), with 999 and 9999 permutations used, respectively. We tested for cophylogeny with the Procrustes Application to Cophylogenetic Analysis (PACo) and ParaFit, implemented in the PACo (Hutchinson et al. 2017) and APE (Paradis, Claude, and Strimmer 2004) R packages, respectively. For both tests, the Cailliez correction (Cailliez 1983) for negative eigenvalues was applied, and 999 permutations were used to assess significance. Tests of trait associations were performed with phylogenetic generalized least squares (PGLS) and randomization of residuals in a permutation procedure

(RRPP), implemented in the phytools and RRPP packages (Collyer and Adams 2018), respectively. To ascertain significance, 999 permutations were used for both methods.

Ancestral state reconstruction models were fit to archaeal taxon abundances (extant traits) via the phylopars method as implemented in the Rphylopars package (Goolsby, Bruggeman, and Ané 2017). The method incorporates intra-species trait variation, so all samples were used instead of employing an intra-species sensitivity analysis (see above). We first compared log-likelihoods of four different models: Brownian Motion, Ornstein-Uhlenbeck, Early-Burst, and Star-Phylogeny. Brownian Motion and Ornstein-Uhlenbeck models had the best log-likelihoods for class- and genus-level archaeal abundances, respectively. Predicted trait values were visualized on the host phylogeny via the Phytools R package.

Tables S6 and S7 list published body temperature and methane emission data used in this study.

Significant patterns of Archaea-Archaea and Archaea-Bacteria co-occurrence were inferred via the cooccur R package (Griffith, Veech, and Marsh 2016). Subnetworks in each co-occurrence network were identified with the walktrap algorithm (Pons and Latapy 2005).

General data manipulation and visualization was performed in R (R Core Team 2020) with the following R packages: dplyr, tidyr, and ggplot2 (Wickham 2009). Phylogenies were manipulated and visualized with the APE and phytools R packages and with iTOL (Letunic and Bork 2016). Networks were manipulated and visualized with the igraph (Csardi and Nepusz 2006), tidygraph (Pedersen 2018b), and ggraph (Pedersen 2018a) R packages. High performance computing cluster job submission was performed via the batchtools (Lang, Bischl, and Surmann 2017) and clustermq (Schubert 2019) R packages. For ASV-specific tests (*e.g.,* LIPA, PGLS, and co-occurrence), only ASVs present in >5% of samples were included. Multiple hypothesis testing was corrected via the Benjamini-Hochberg procedure.

### Supplemental Results

*Prevalence and diversity of Archaea across vertebrate clades*

Of 311 genomic DNA samples from 5 vertebrate taxonomic classes, 185 (60%) passed 16S rRNA PCR amplification, MiSeq sequencing, and sequence data quality control (Table S2). Success rates were highest for Reptilia (73%) and Aves (67%), 58% for Mammalia, 50% for Amphibia, and 50% for Reptilia (Figure S2). The 185 successful samples comprised mostly wild individuals (76%) and a total of 110 species, with a mean 1.7 ± 4.3 s.d. samples per species (Figure S1). Mammalia made up the majority of samples (72%); still, non-mammalian samples spanned 22 families and 35 genera. In regards to diet, success rates were 70, 56, and 46% for herbivores, omnivores, and carnivores, respectively (Figure S2). Feces samples had a substantially higher success rate (62%) versus gut contents (38%), but there was little difference between wild and captive individuals (62 versus 56%, respectively). The mean per-species success rate was 61% ± 49 s.d., and when just assessing species with >1 sample (72 of 158), the success rate was 63% ± 37.3 s.d. Plotting the number of successful and failed samples onto a phylogeny of all species showed that success often varied among individuals of a species (Figure S3). In addition, some phylogenetic clustering of success rates could be observed. Indeed, when just considering mammalia, which made up the majority of samples

(73%), the orders Lagomorpha, Carnivora, and Rodentia had the lowest success rates (<50% for each), while success rates were 100% for Monotremata, Perissodactyla, and Proboscidea (Figure S2). While these findings are compelling, one must consider that failure may have resulted from many phenomena besides absence of Archaea from the gut, such as PCR inhibitors or insufficient DNA for effective amplification. Still, success across highly varied host taxonomic groups, diets, and sample types indicates that Archaea are widespread among vertebrates, regardless of diet.

Rarefaction analysis using the Shannon index revealed that archaeal diversity saturated at a low sampling depth of approximately 250 sequences, regardless of the host class (Figure S4). We confirmed these results with another rarefaction method that extrapolates diversity beyond obtained sampling depths, with diversity based on Hill numbers (Figure S4). These results contrast most gut microbiome studies using the commonly used “universal” Earth Microbiome 16S rRNA primer set 515F-806R, in which bacterial and archaeal diversity is usually not saturated for the sampling depths reached (Walters et al. 2016; Thompson et al. 2017; Youngblut et al. 2019).

The dataset comprised 1891 amplicon sequence variants (ASVs), with a rather diverse taxonomic composition for Archaea, comprising 6 phyla (Asgardaeota, Crenarchaeota, Diapherotrites, Euryarchaeota, Nanoarchaeaeota, Thaumarchaeota) and 10 classes (Figure 1). Class-level taxonomic compositions were fairly consistent among individuals of each host species (Figure S5; Table S3). We note that Asgardarchaeota and Diapherotrites were each only represented by 1 ASV, and each were found in only 1 species: Asgardaeota in the the European Otter (*Lutra lutra*) and Diapherotrites in the Smooth Newt (*Lissotriton vulgaris*). Neither clade is known to be animal-associated (Borrel et al. 2020). Also, the Thermococci class (Euryarchaeota phylum) comprised only 2 ASVs, with one only found in the Common Carp (*Cyprinus carpio*) and the other in the European Otter. Both ASVs were classified as Methanofastidiosales, with one identified as Methanofastidiosum. No member of this class is known to be host-associated (Söllinger and Urich 2019; Borrel et al. 2020). Plotting mean abundances of taxonomic classes onto a tree of all species revealed that Methanobacteria (Euryarchaeota phylum) dominated in many species, but dramatically different microbiome compositions were observed scattered across the phylogeny. For instance, Thermoplasmata (Euryarchaeota phylum) dominated in multiple non-human primates, while two Mammalia and one Aves species were nearly completely comprised of Nitrososphaeria (Thaumarchaeaota phylum): the European badger (*Meles meles*), the Western European Hedgehog (*Erinaceus europaeus*), and the Rook (*Corvus frugilegus*). Halobacteria (Euryarchaoeota phylum) dominated the Goose (*Anser anser*) microbiome, which were all sampled from salt marshes. The class was also noticeably present in some distantly related animals inhabiting high salinity biomes (*e.g.,* the Nile Crocodile and the Short Beaked Echidna; Tables S1 & S3). Bathyarchaeia, a class in the Crenarchaea phylum according to the SILVA database taxonomy but also known as the Candidatus Bathyarchaeota phylum, are not known to inhabit the vertebrate gut (Borrel et al. 2020); however, we observed a total of 9 Bathyarchaeia ASVs in 8 samples, comprising 6 species spanning 4 taxonomic classes (all except Mammalia; Table S4). The total relative abundance was <0.5% in 4 of the species, while substantially higher (3.3%) in the Nile Crocodile (*Crocodylus niloticus*), and quite abundant in 2 Smooth Newt samples (17.9 and 42.2%).

Only 40% of ASVs had a ≥97% sequence identity match (a pseudo-species level) to any cultured representative in the All Species Living Tree database (Figure S6A). Of the 10 taxonomic classes represented by all ASVs, 5 had no match at ≥85% sequence identity: Odinarchaeia, Bathyarchaeia, Iainarchaeia, Woesarchaeia, and Thermococci. Taxonomic novelty to cultured representatives differed substantially among the other 5 classes (Figure S6B); only Methanobacteria had >50% ASVs with a species-level match (52%), while <20% of ASVs belonging to Thermoplasmata and Nitrososphaeria had such a match. These findings suggest that our dataset consists of a great deal of uncultured taxonomic diversity.

#### *Archaea-targeting primers reveal much greater archaeal diversity*

We compared the archaeal diversity identified with the archaeal-targeting primers ("16S-arc") used in this study to the standard "universal" 16S rRNA primers ("16S-uni") used by Youngblut and colleagues on many of the same samples (Youngblut et al. 2019). Importantly, both datasets were processed in the same manner (see Methods). A total of 140 samples overlapped between the two datasets, with the majority of species (77%) consisting of mammals, but all 5 classes were represented (Figure S7). The 16S-uni primers generated a total of 169 ASVs, which is only 12.1% of archaeal ASVs generated by the 16S-arc primers for the same samples. All archaeal classes except the Soil Crenarchaeal Group were substantially more represented in the 16S-arc dataset, with 6 classes completely absent from the 16S-uni dataset: Nitrososphaeria, Woesarchaeia, unclassified Eukyarchaeota, Iainarchaeia, Bathyarchaeia, and Odinarchaeia. Besides the Soil Crenarchaeal Group, class-level prevalence across host species was substantially higher across hosts when grouped by taxonomic class or diet (Figure S7). For example, Methanobacteria was observed in all host species via the 16S-arc primers, while prevalence dropped substantially for 16S-uni primers (*e.g.,* only 9% for Aves). These findings show that the "universal" NGS 16S rRNA primers used for most microbiome studies can substantially undersample archaeal diversity, as previously observed (Raymann et al. 2017; Koskinen et al. 2017; Pausan et al. 2019)

#### *Host diet and evolutionary history explain various aspects of archaeal diversity*

We used multiple regression on matrices (MRM) to assess which potential factors explain archaeal beta diversity. We employed this approach because archaeal beta diversity, host phylogenetic relatedness, and geographic distance can be inherently represented as distance matrices, while distances can be calculated for other explanatory factors such as similarity of detailed diet compositions (see Methods). Due to a lack of within-species phylogenetic relatedness data, we used one individual per host species and assessed intra-species variation by repeating the analysis 99 more times, each time with one randomly selected individual per species. Unless otherwise noted, this permutation-based intra-species sensitivity analysis was used for all hypothesis testing.

Geographic distance, habitat, and technical components (*e.g.,* feces versus gut contents) did not significantly explain beta diversity, regardless of the diversity metric (Figure 2A). Host phylogeny significantly explained diversity as measured by unweighted UniFrac, Bray Curtis, and Jaccard; however, significance was not quite reached for weighted UniFrac. The percent variation explained was dependent on the beta diversity measure and varied from ~28%

for Jaccard to ~12% for unweighted UniFrac. In contrast to host phylogeny, diet was only explanatory for Bray-Curtis, with ~12% of variance explained. Mapping the major factors onto ordinations qualitatively supported our results (Figure S8). Applying the same MRM analysis to just non-mammalian species did not generate any significant associations between host phylogeny or diet (Figure S9), likely due to the low sample sizes (*n* = 39). However, host phylogeny did have comparable coefficients as when including all species and were nearly significant for both the Bray-Curtis and Jaccard indices, while diet showed no such trend towards significance. These findings suggest that host evolutionary history mediates vertebrate gut archaeal diversity more than diet, with diet mainly altering the abundances of archaeal ASVs shared by various hosts, while host phylogeny also alters the composition of archaeal taxa.

We also assessed alpha diversity via MRM in order to provide a consistent comparison to our beta diversity assessment, with alpha diversity represented here as a euclidean distance matrix (Figure S10). In contrast to beta diversity, no factors significantly explained alpha diversity calculated via either the Shannon Index or Faith's Phylogenetic Diversity (Faith's PD). Of note, geographic distance nearly significantly explained Shannon Index diversity (P = 0.06), while the same was true of habitat for Faith's PD (P = 0.16).

*A signal of Archaea-Vertebrata co-phylogeny*

To test for corresponding phylogenetic associations on both the host phylogeny and the archaeal 16S rRNA phylogeny, we employed two approaches to quantify signals of co-phylogeny: Procrustes Application to Cophylogenetic Analysis (PACo) and ParaFit (Paradis, Claude, and Strimmer 2004; Hutchinson et al. 2017). Both PACo and ParaFit tests were both significant (P < 0.01) for each of the 100 permutations of subsampling one individual per host species, indicating a signal of co-phylogeny that is robust to intra-species microbiome variation. We investigated which host species showed the strongest signal of cophylogeny by assessing the distribution of PACo Procrustes residuals, which provide an indication of local congruence between phylogenies (lower residuals indicate a stronger congruence). Mammalia showed a substantially stronger association relative to the other four classes (Figure 2D), with residuals decreasing in the order of Actinopterygii > Amphibia > Reptilia > Aves ≫ Mammalia, and these differences were significant (Kruskal-Wallis < 0.01; pairwise Wilcox < 0.01 for all). In regards to diet, residuals were significantly lower for herbivores relative to omnivores and carnivores (Wilcox, P < 0.0001), while carnivores and omnivores did not significantly differ (Figure 2E).

*Specific archaeal ASVs are associated with host phylogeny*

Given the evidence of host phylogeny explaining aspects of archaeal gut microbiome diversity, we sought to further resolve this association by testing whether archaeal taxon abundance is clustered on the host phylogeny. We found 37 ASVs to show significant global phylogenetic signal (Pagel's $\lambda$, adj. P < 0.05) spanning three phyla: Euryarchaota, Thaumarchaeota, and Crenarchaeota (Figure 2C). The clade with the highest number of significant ASVs (*n* = 15) was Methanobacteriaceae, followed by Nitrososphaeraceae (*n* = 12), and Methanocorpusculaceae (*n* = 5). While lambda coefficients varied across ASVs, most showed a very strong association (Pagel's $\lambda$ > 0.9), with major exceptions being a Methanosarcinaceae ASVs and an unclassified Methanomicrobia ASV (Figure 2C).

We next tested for local phylogenetic signals to resolve archaeal taxon specificities for particular host clades. We used the local indicator of phylogenetic association (LIPA) and found 25 ASVs to have significant associations with certain host clades. Mapping significant associations on the host phylogeny revealed that clade-specificity was generally shallow and often spanned only 2 species (Figure S12). For instance, 4 Nitrososphaeraceae ASVs were associated with 2 snake species (*Zamenis longissimus* and *Natrix natrix*), 3 Methanobrevibacter ASVs were associated with 2 species of kangaroo (*Macropus giganeus* and *Macropus fuliginosus*), and a Methanocorpusculum ASV was associated with both camel species (*Camelus dromedarius* and *Camelus bactrianus*). The 2 major exceptions to this trend were the Methanothermobacter ASVs, which associated with many species of Aves, while the Methanobrevibacter and Methanosphaera ASVs associated with many Artiodactyla species (true ruminants; Figures S12). Summarizing the number microbe-host clade associations revealed clear partitioning of archaeal taxa by host clade, except for Methanobrevibacter, for which at least one ASV was associated with each host order for which any phylogenetic signal was observed (*n* = 23; Figure S12B). Altogether, these results help to resolve which particular archaeal clades are most strongly associated with host evolutionary history.

We also tested for phylogenetic signal of alpha diversity but found no significant global associations when measuring diversity via the Shannon Index or Faith's PD ($P > 0.05$) and no local associations (adj. $P > 0.05$). These findings correspond with our MRM analysis of alpha diversity in that host phylogenetic relatedness does not seem to correspond with total archaeal diversity in the gut.

*Specific methanogen ASVs are associated with diet*

We used two methods to resolve the specific effects of diet on the archaeal microbiome while controlling for host evolutionary history: phylogenetic generalized least squares (PGLS) and randomization of residuals in a permutation procedure (RRPP). The former is a common test for association between traits while controlling for phylogenetic relatedness, while the latter can exhibit higher statistical power while minimizing false positives (Revell 2010; Collyer and Adams 2018). PLGS identified 10 ASVs as being significantly associated with diet (adj. $P < 0.05$; Figure S11). All ASVs belonged to the Euryarchaeota phylum, and comprised 4 genera: Methanobrevibacter, Methanosphaera, Methanothermobacter, and candidatus Methanomethylophilus. The RRPP analysis identified the same 10 ASVs along with 5 more that belonged to the same genera (Figure 2B). We used the RRPP models to predict ASV abundances with 95% confidence intervals (CIs) for each diet in order to determine diet-specific enrichment. Methanobacteria ASVs differed in their responses to diet, with 5 being most abundant in herbivores, while the other 6 were more abundant in omnivores/carnivores (Figure 2B). Notably, diet enrichment differed even among ASVs belonging to the same genus. In contrast to the Methanobacteria ASVs, all 4 Methanomethylophilus ASVs were predicted as more abundant in omnivores/carnivores. These findings suggest that diet influences the abundances of particular ASVs, and even closely related ASVs can have contrasting associations to diet. All significant ASVs were methanogens, which may be due to the species studied (e*.g.,* a mammalian bias) or possibly because certain methanogens respond readily to diet, possibly due to syntrophic associations with diet-specific bacteria.

When applied to alpha or beta diversity, neither PGLS nor RRPP identified any significant associations with diet after accounting for host phylogenetic relatedness. These findings correspond with our MRM analyses by indicating that diet is not a strong moduator of overall archaeal diversity in the vertebrate gut, although certain ASVs do seem to be substantially affected (Figures 2B & S11).

*Evidence of widespread Methanobacteria presence in the ancestral vertebrate gut*

We utilized ancestral state reconstruction (ASR) to investigate which archaeal clades were likely present in the ancestral vertebrate gut. Traits were defined as archaeal taxon abundances. Notably, we used a method that incorporated intra-species trait variance, allowing us to directly utilize the entire host dataset for the reconstruction (see Methods). Our model for predicting class-level abundances was overall quite accurate at extant species trait prediction (adj. $R^2$ = 0.86, P < 2e-16; Figure S14). However, predictions were not accurate for 2 of the 6 classes (Halobacteria and Nitrososphaeria, P > 0.1), likely due to low prevalence across extant host species (Figures 2 & S15). Excluding the poorly predicted classes, the 95% CIs for predicted abundances were constrained enough to be informative (mean of 26 % ± 29 s.d.) across extant and ancestral host species. The model revealed that Methanobacteria was uniquely pervasive across ancestral nodes, while other classes were sparsely distributed among extant taxa and across a few, more recent ancestral nodes (Figures 2 & S15). Moreover, the model predicted that Methanobacteria was the only class to be present in the last common ancestor (LCA) of all mammals and the LCA of all 5 host taxonomic classes (Figure 3B & 3C).

We also generated an ASR model for genus-level abundances of all genera in the Methanobacteria class in order to resolve the association between Methanobacteria clades and the ancestral vertebrate gut. Our model was somewhat more accurate at predicting extant traits than our class-level model ($R^2$ = 0.93, P < 2e-16; Figure S14), and all 4 genera were accurately predicted (P < 5.5e-10 for all). Predicted trait value 95% CIs were again informative (mean of 28 ± 24 s.d.). The model predicted 3 of the 4 genera to be present in the LCA of all mammals and the LCA of all host species (Figure 3F & 3G). Of the 3, Methanobrevibacter and Methanothermobacter were predicted to have similar abundances for both LCAs (~30-35%), while Methanosphaera was much lower (~5%). Mapping predicted abundances onto the host phylogeny revealed that Methanobrevibacter was predicted as most highly abundant in the Artiodactyla and generally abundant across most Mammalia clades (Figure S16). In contrast, Methanothermobacter was predicted to be most highly abundant and prevalent across the Aves and also mammalian clades in which Methanobrevibacter was less abundant (*e.g.,* Carnivora and Rodentia). Methanosphaera was predicted to be prevalent across most animal clades, but generally at low abundance.

*Methanothermobacter abundance is correlated with body temperature*

Methanothermobacter is not known to be host-associated (Borrel et al. 2020); still, we observed a total of 39 Methanothermobacter ASVs spanning 78 samples (mean of 18 ± 30 s.d. samples per ASV), which strongly suggests that its presence is not due to contamination. Moreover, the top BLASTn hit for 36 of the 39 ASVs was to a cultured Methanothermobacter strain (Figure S17, Table S5), including the top 15 most abundant ASVs, which indicates that the taxonomic annotations are demonstrably correct.

The high prevalence of Methanothermobacter among Aves lead us to the hypothesis that body temperature significantly affects the distribution Methanothermobacter (Figure S18), given that birds generally have higher body temperatures than mammals (Clarke and O'Connor 2014) and all existing Methanothermobacter cultures are thermophiles (Bonin and Boone 2006). Moreover, Methanothermobacter is not abundant in Monotremata and Marsupialia species relative to the placental groups, which reflects a lower body temperature in the latter clades (Figure S18). We were able to assign published body temperature data to 73 mammalian and avian species (Figure S19A & S19B; Table S6). Genus-level abundances of Methanothermobacter significantly correlated with body temperature (RRPP, adj. $P < 0.001$), while Methanobrevibacter and Methanosphaera did not (Figures S19C & S19D). However, the association was only significant if not accounting for host phylogeny (RRPP, adj. $P > 0.05$), indicating that the association between Methanothermobacter and body temperature could not be decoupled from host evolutionary history. We also identified 7 Methanothermobacter ASVs to be correlated with body temperature (RRPP, adj. $P < 0.05$; Figure S19E), while no Methanobrevibacter or Methanosphaera ASVs were correlated. Again, the association was only significant if not accounting for host phylogeny. Regardless, we provide evidence congruent with the hypothesis that Methanothermobacter abundance is modulated by host body temperature and is thus rather highly abundant in birds and various placental mammal clades.

We note that among the host species in which methane emission data exists (Hackstein and van Alen 1996; Clauss et al. 2020), avian species with high abundances of Methanothermobacter have emission rates on the higher end of mammal emission rates (Figure S20), suggesting that Methanothermobacter is indeed a persistent inhabitant in the gut of some avian species.

*Microbe-microbe interactions modulating archaeal diversity*

Besides host-specific factors potentially modulating diversity, microbe-microbe interactions may also play a significant role. We first tested for solely archaeal interactions by inferring instances of co-occurrence among archaeal ASVs. The co-occurrence network contained clearly defined subnetworks, with few significant positive associations between them (Figure S22), especially for the largest 6 subnetworks (Figure S21). The only significant negative co-occurrences were between Subnetwork 1, which was dominated by Methanobrevibacter, and Subnetwork 4, which was dominated by Methanothermobacter. These 2 subnetworks differed substantially in their distributions across host clades, with Subnetwork 1 ASVs only highly prevalent among Artiodactyla, while Subnetwork 4 ASVs were highly prevalent across a number of mammalian orders (*e.g.,* Carnivora and Rodentia) and almost all avian orders (Figure S23). Among subnetworks, ASV taxonomy was highly homogeneous. Indeed, we found ASVs to significantly and strongly associated with those of the same clade versus from other clades, regardless of taxonomic level (Figure S21C), although assortativity by taxonomic affiliation substantially dropped between the family and genus levels.

We investigated potential diet-specific archaea-archaea interactions by separately testing for co-occurrences across samples of each diet (Figure S24). The number of significant co-occurrences dropped from herbivores ($n$ = 560) to omnivores ($n$ = 134) to carnivores ($n$ =

81). In contrast, assortativity by taxonomic group was generally lowest for omnivores and highest for carnivores, regardless of taxonomic level. These findings suggest that the carnivore gut is composed of simpler and more taxonomically homogenous archaeal consortia relative to omnivores and herbivores.

We also assessed Bacteria-Archaea interactions by utilizing the overlapping 16S-uni dataset samples from Youngblut and colleagues (Youngblut et al. 2019). Prior to merging the datasets, we removed all archaeal ASVs from the 16S-uni dataset. Archaeal and bacterial alpha diversity were not correlated, regardless of measuring diversity via the Shannon Index or Faith's PD (Pearson, $P > 0.05$; Figure 4). Moreover, archaeal and bacterial beta diversity were not correlated (Mantel, $P > 0.05$; Procrustes superimposition, $P > 0.05$), regardless of the measure: Bray-Curtis, Jaccard, and weighted/unweighted UniFrac. These results suggest that archaeal diversity is not explained by bacterial diversity nor vice versa.

Inferring a co-occurrence network of bacterial and archaeal ASVs revealed a large number of significant co-occurrences ($n$ = 3018; Figure 4); all of which were positive. Bacteria-Archaea and Archaea-Archaea associations comprised 13.1 and 6.1% of the network edges, respectively. While overall network taxonomic assortativity was low, assortativity of just Archaea was quite high (≥0.774 for all taxonomic levels). The entire network comprised 5 subnetworks, but only 2 included archaea: one of which included only Methanobrevibacter ASVs, while the other was dominated by Methanothermobacter ASVs. The Methanobrevibacter-only subnetwork also comprised 13 bacterial families from 3 phyla. Firmicutes dominated among the bacterial ASVs (87%), with Bacteroidetes as a distant second (11%). The most represented bacterial families in the network were Ruminococcaceae (46%), Lachnospiraceae (13%), and Christensenellaceae (11%), which are known include hydrogen generating species that often occur with Methanobrevibacter (Hansen et al. 2011; Goodrich et al. 2014; Borrel et al. 2020). The Methanothermobacter-dominated subnetwork included much less bacterial diversity, with only 3 families: Burkholderiaceae (Proteobacteria phylum); Enterococcaceae and Clostridiaceae 1 (Firmicutes phylum). These findings indicate that a subset of archaeal ASVs co-occur with specific bacterial ASVs in each of the 2 consortia: the Methanothermobacter-dominanted consortium most prevalent among birds and the Methanobrevibacter-dominated consortium most prevalent among ruminants and various other plant-consuming mammals (Figure S18). While only methanogens were observed to co-occur with bacteria, this may be due to the mammalian bias of the dataset, given that prevalence of non-methanogenic archaea is lower among mammals relative to other vertebrate classes (Figure 1).

### Supplemental Tables

**Table S1.** All relevant metadata for all samples in the 16S rRNA amplicon dataset.

**Table S2.** Metadata for all samples in which Archaea-targeted 16S rRNA amplicon library preparation and sequencing was attempted (*n* = 311) and the samples that passed all quality control measures (*n* = 185).

**Table S3.** Percent relative abundance of each archaeal taxonomic class in each sample (*n* = 185). Classes are labeled as “Phylum;Class”.

**Table S4.** Genus-level percent relative abundances of Bathyarchaeia in all samples where the clade was detected.

**Table S5.** The top 5 BLASTn hits of all Methanothermobacter ASV sequences to the All Species Living Tree dataset (see Methods). Mean percent relative abundances across all samples and samples grouped by host taxonomic class are also provided.

**Table S6.** Publicly available body temperature data used in this study. If multiple temperature data points per species were available, the mean temperature was used. The datasets include “Clarke2010” (Clarke, Rothery, and Isaac 2010), “Clarke2014” (Clarke and O’Connor 2014), “McNab1966” (McNab 1966), “Prinzinger1991” (Prinzinger, Preßmar, and Schleucher 1991), “Riek2013” (Riek and Geiser 2013), “Sieg2009” (Sieg et al. 2009), and “Teare2002” (Teare 2002). “No match” indicates the species lacking a match to any of the body temperature datasets; these species were not included in any analyses of body temperature due to a lack of data.

**Table S7.** Publicly available animal methane emission data used in this study. The studies comprise “Hackstein_1996” (Hackstein and van Alen 1996) and “Clauss_2020” (Clauss et al. 2020).

### Supplemental Figures

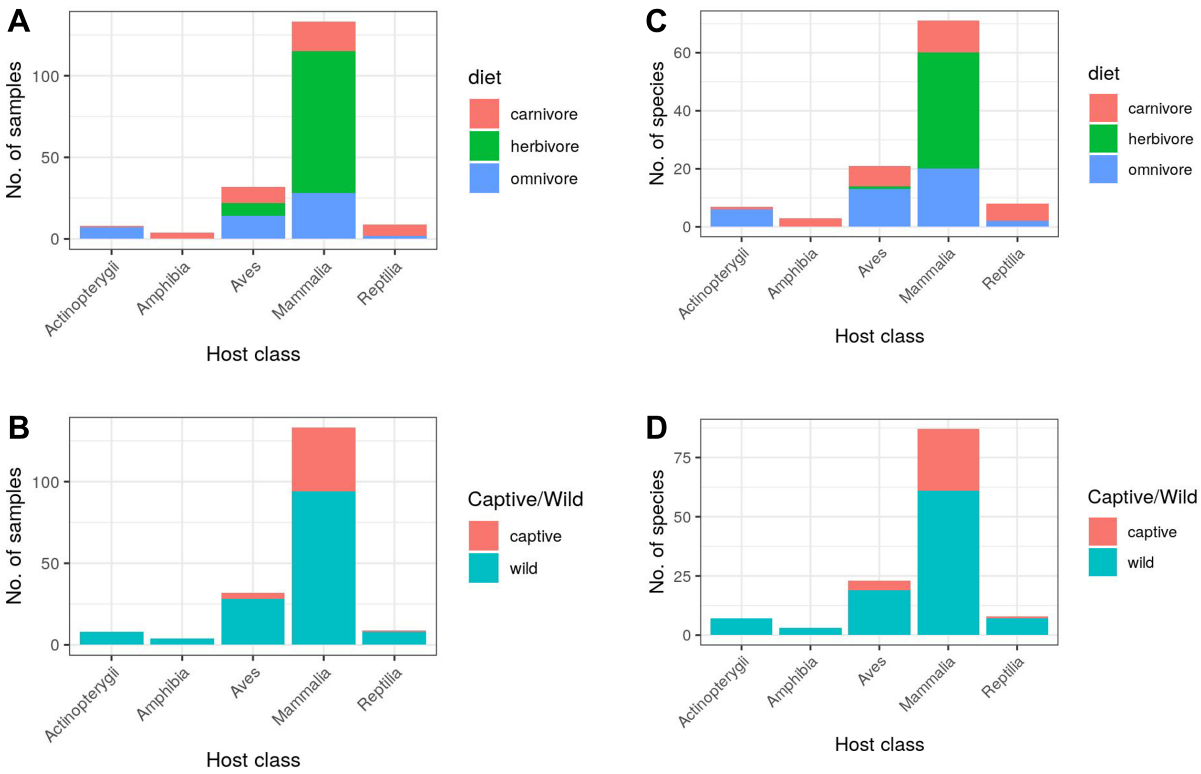


**Figure S1.** The number of samples (A & B) or host species (C & D) in the final sequence dataset, grouped by host class, host diet (A & C) or host captive/wild status (B & D).

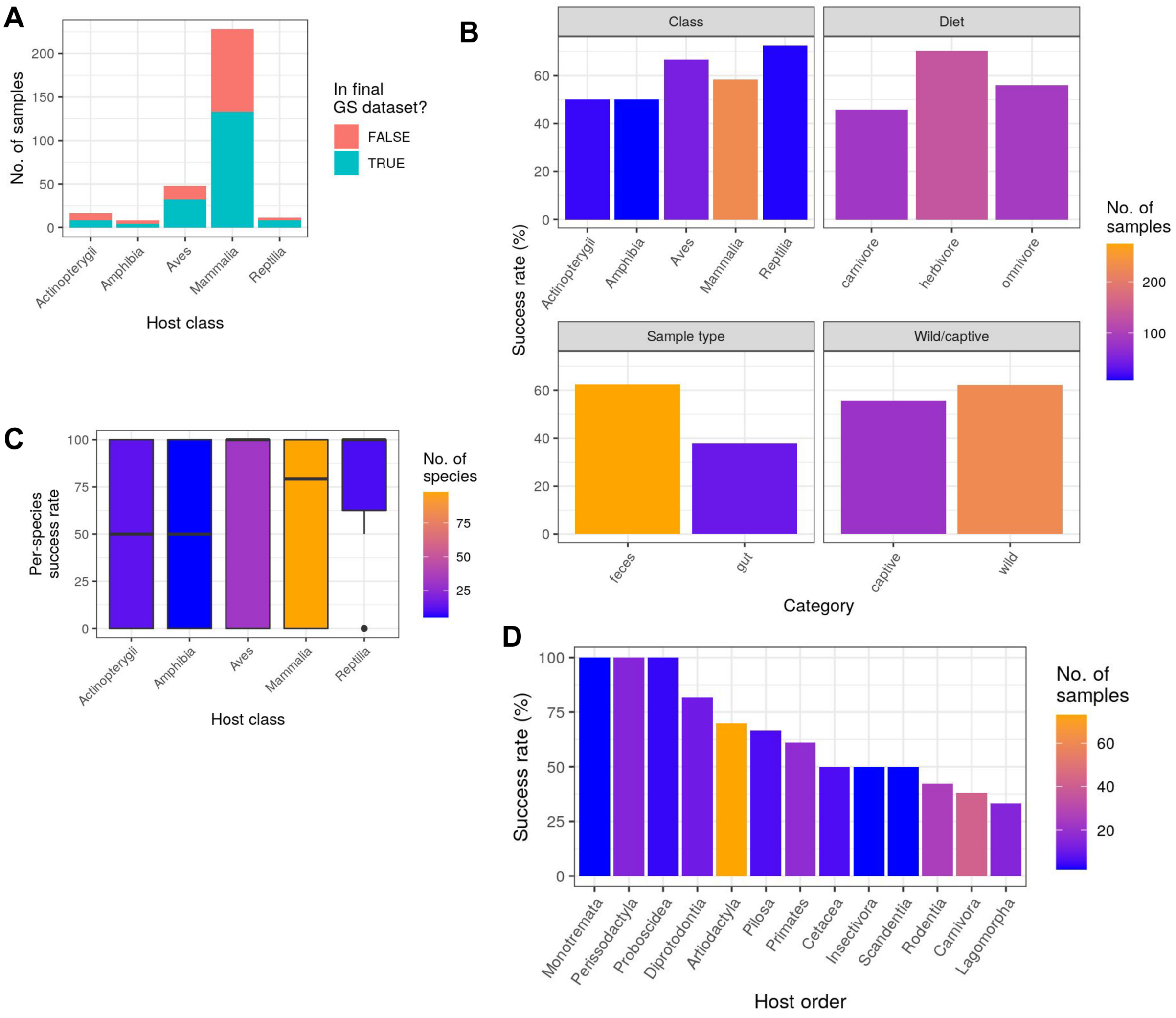


**Figure S2.** A) The number of samples that passed or failed PCR amplification and sequence data quality control. B) The percent of total samples that passed PCR amplification and sequence data quality control (*i.e.,* the success rate), with values grouped by various host metadata categories. C) The success rate among individuals of the same species, grouped by host class. D) The success rate for each mammalian taxonomic order. See Table S2 for a list of all successes and failures.

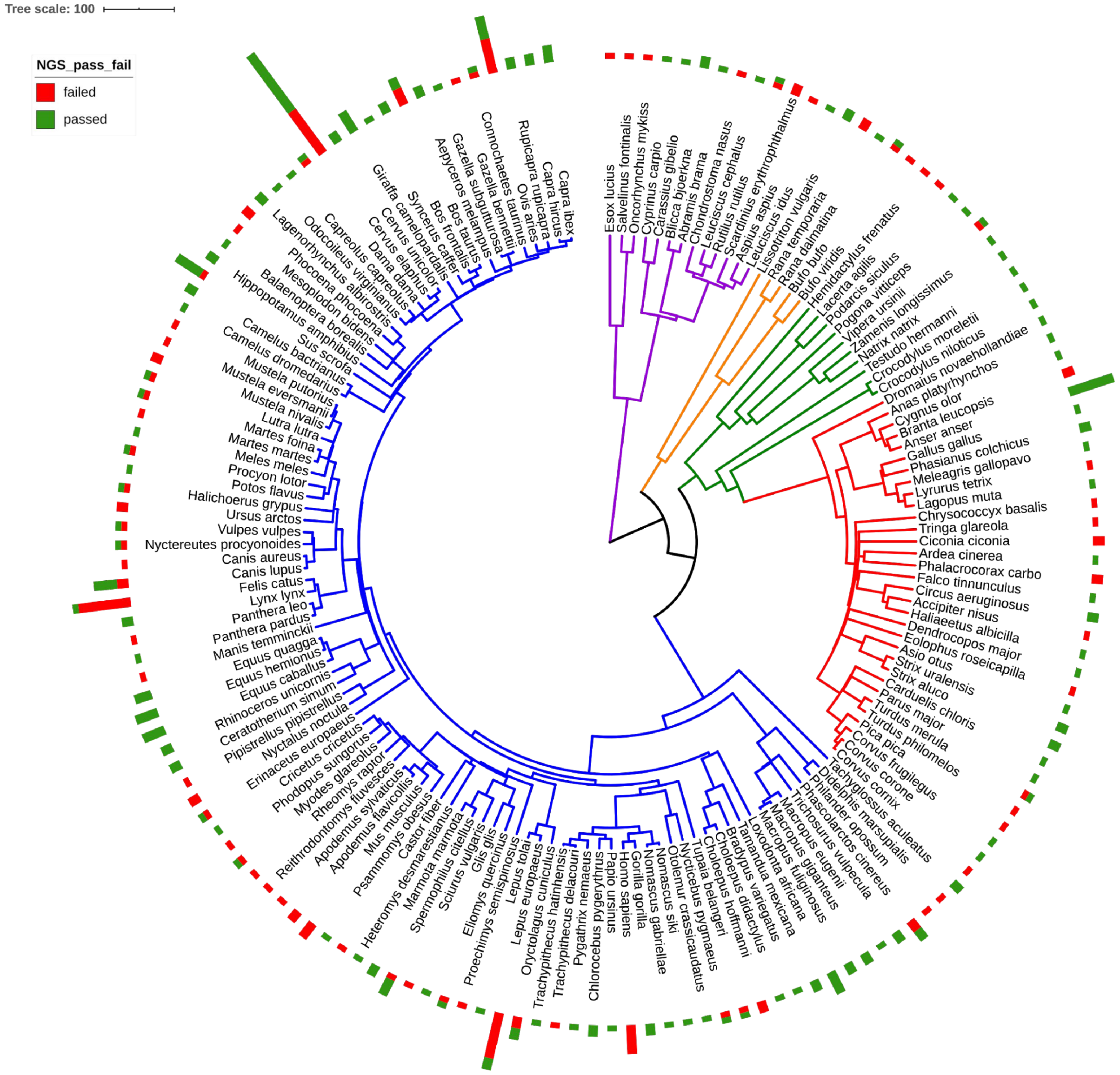


**Figure S3.** The number of samples that passed PCR amplification and sequence data quality control ("passed") and those that failed ("failed") mapped onto a phylogeny of all host species. The phylogeny is the same as shown in Figure 1.

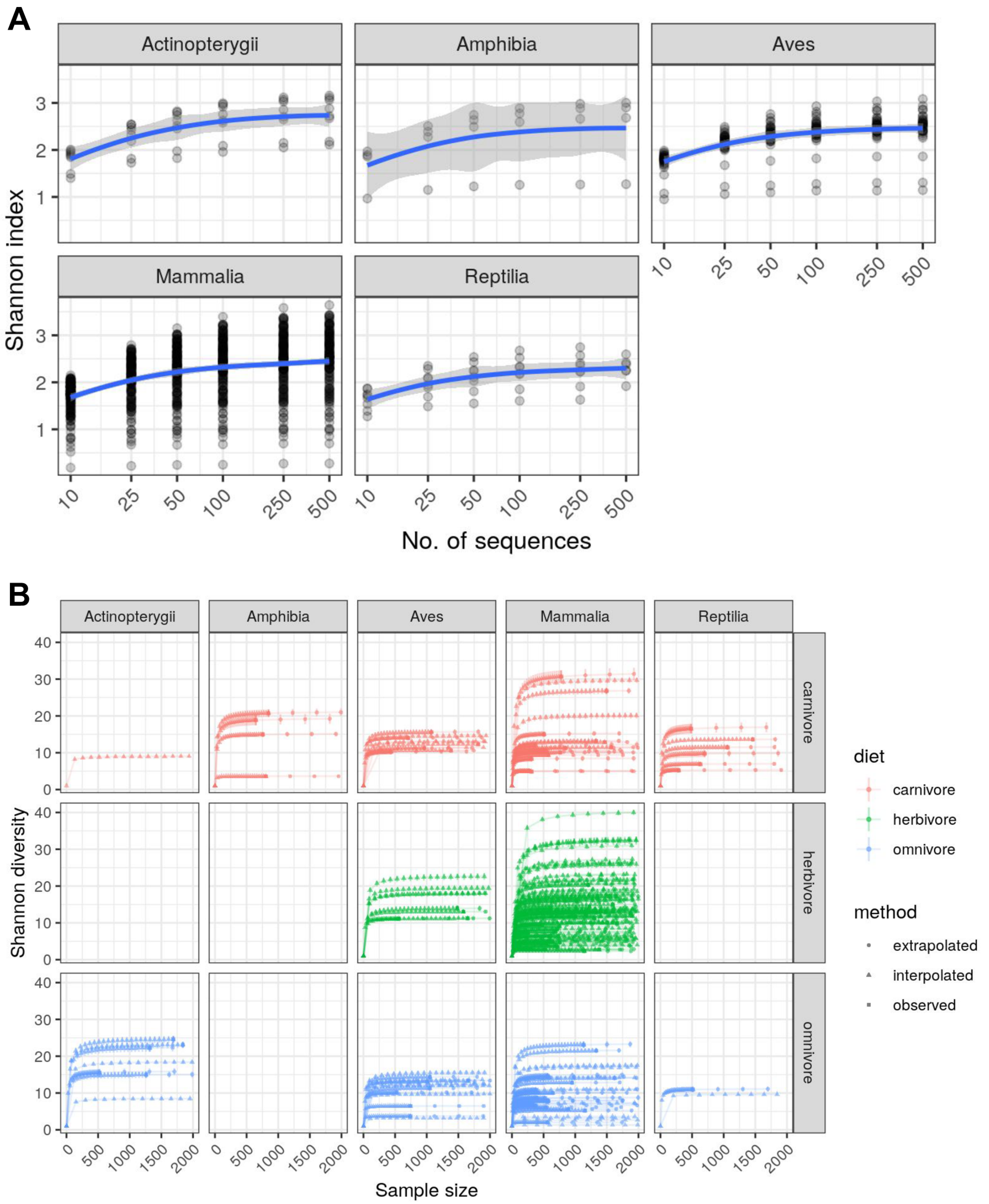


**Figure S4.** A) Rarefaction grouped by host taxonomic class, with subsampling continued up to 500 per sample (if possible, depending on the sample). The blue lines are a smoothed curve fit, with grey regions denoting the 95% CI. B) Rarefaction with extrapolation via iNEXT, with subsampling/extrapolation up to 2000 per sample. Diversity was measured as Hill numbers (diversity order of 1, which is equivalent to Shannon diversity).

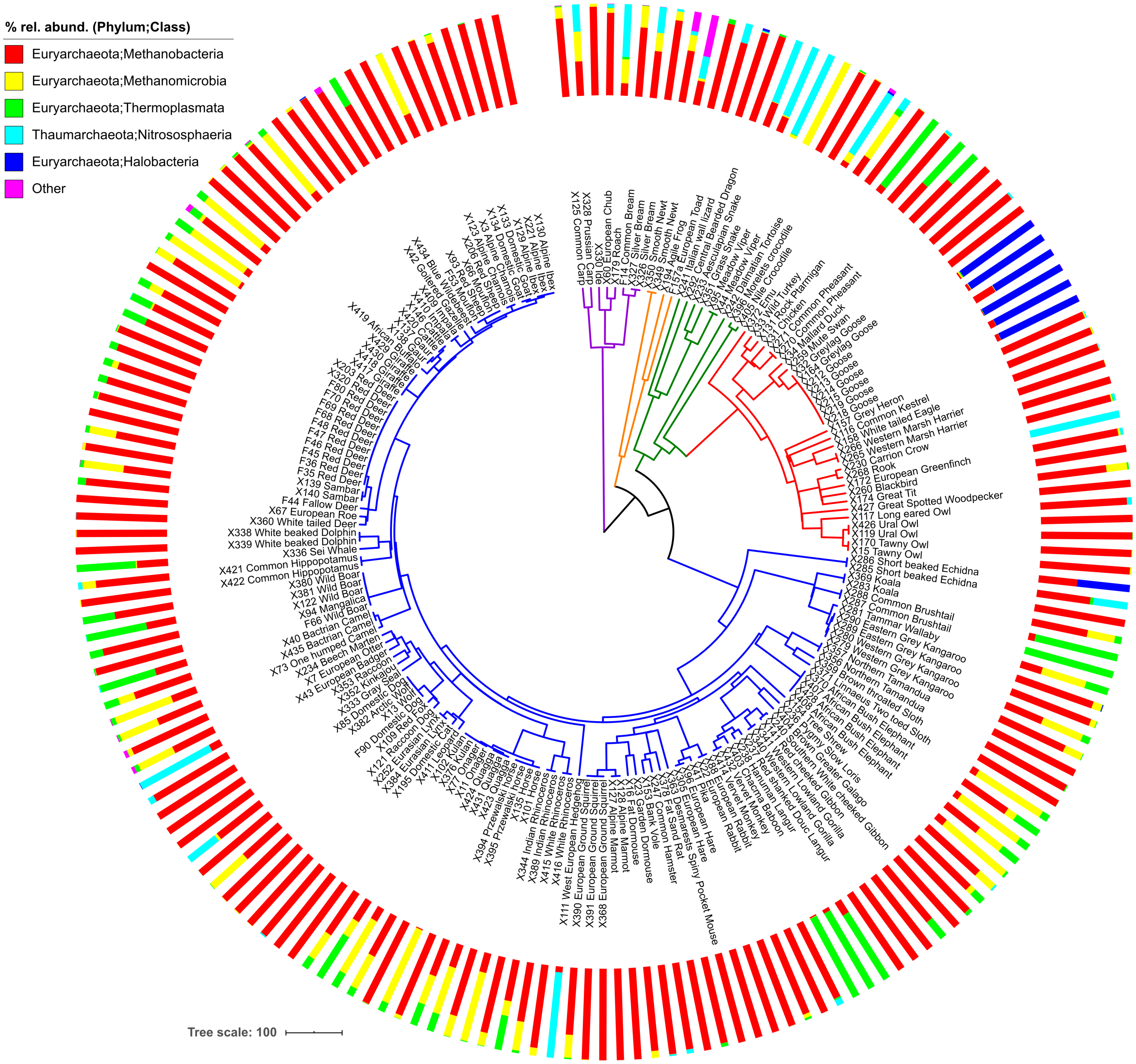


**Figure S5.** The host phylogeny is that same as shown in Figure 1, except tips have been expanded to include all individuals of each species (*n* = 185). Relative abundances of ASVs aggregated by taxonomic class are mapped onto the tree. All classes with <1% mean abundance are labeled as “Other”, which includes Woesearchaeia, Thermococci, Iainarchaeia, and Odinarchaeaeia.

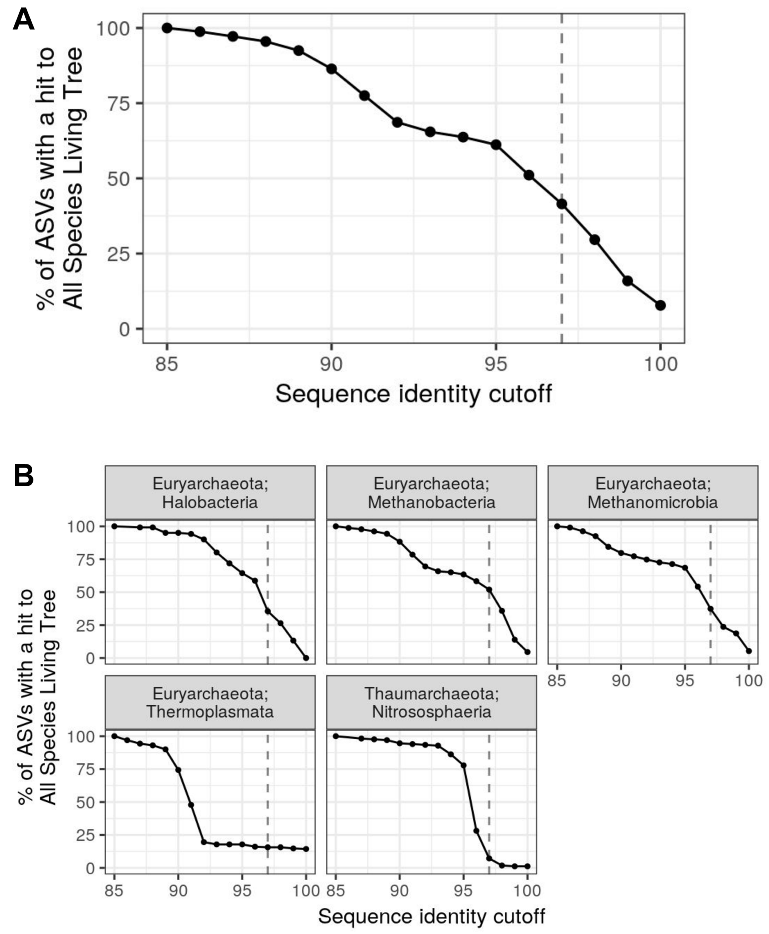


**Figure S6.** *Substantial uncultured archaeal diversity even among relatively well-studied clades*. The percent of ASVs with a ≥1 BLASTn hit to a culture representative in the All Species Living Tree database v132 (hit alignment length ≥95% of the query), depending on the sequence identity cutoff of the BLASTn hit. Values are shown for A) all ASVs and B) ASVs grouped by taxonomic class (facet labels are "Phylum; Class") for the subset of classes in which any hits were observed along the range of sequence identity cutoffs shown.

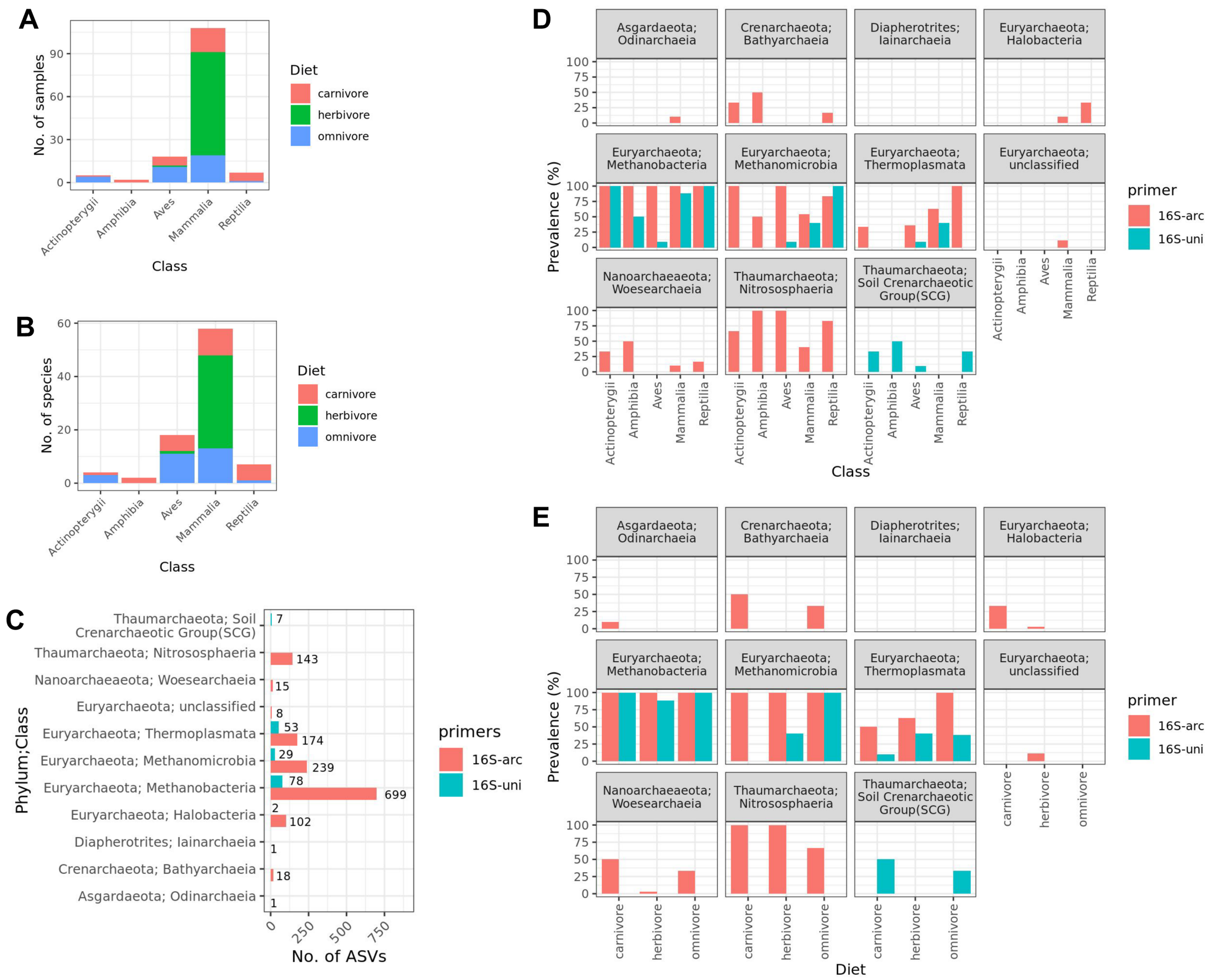


**Figure S7.** *Archaeal-targeting primer set revealed much more archaeal diversity than standard "universal" 16S rRNA NGS primers.* The number of A) samples or B) host species that overlap between the 16S-arc and 16S-uni amplicon sequence datasets. C) The number of archaeal ASVs per sequence dataset. D) & E) The number of archaeal classes across host species grouped by D) host taxonomic class or E) diet.

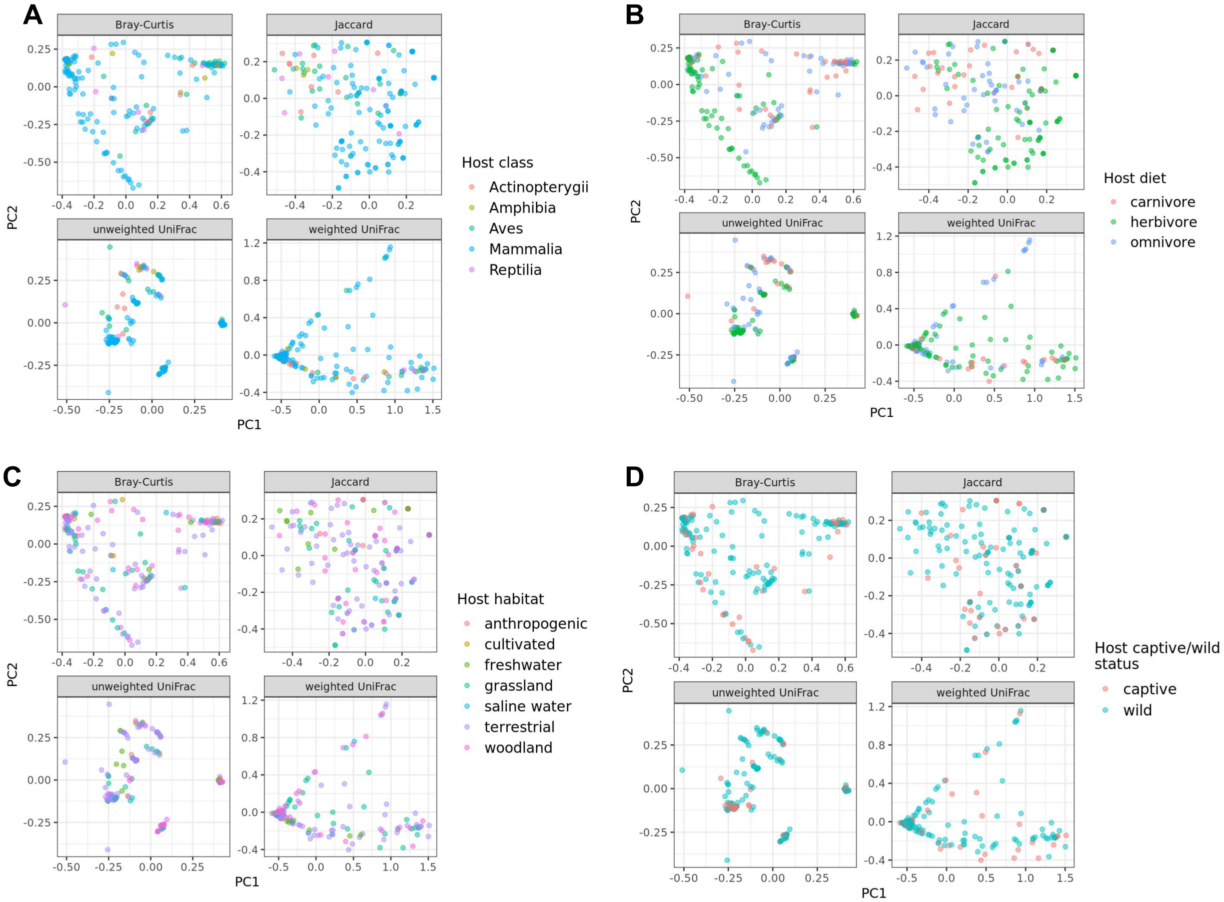


**Figure S8.** *Principal coordinates plots qualitatively agree with the MRM analysis results*. Principal coordinates (PCoA) ordinations of unweighted and weighted Unifrac, Jaccard, and Bray-Curtis distances among all samples, with samples colored by host A) class, B) diet, C) habitat, and D) captive/wild status. The percent variance explained by PC1 and PC2 is 18 & 9 % for Bray-Curtis, 14 and 6 % for Jaccard, 29 and 19 % for unweighted UniFrac, and 72 and 12 % for weighted UniFrac, respectively.

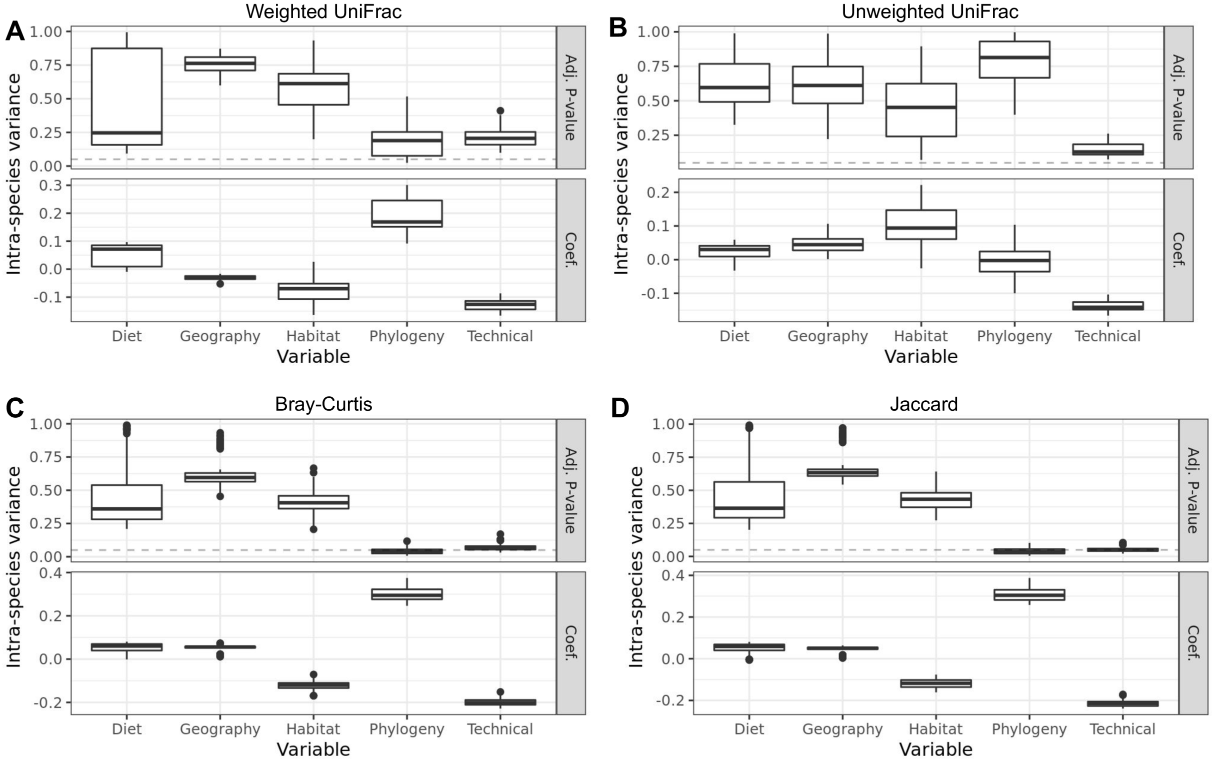


**Figure S9**. *Host phylogeny trending to significance for non-mammalian species.* The plots show the distribution of P-values (“Adj. P-value”) and partial regression coefficients (“Coef.”) across 100 dataset permutations used for multiple regression on matrix (MRM) tests. Unlike Figure 2A, all Mammalia species were excluded, leaving 39 non-mammalian species. For each permutation, one individual per host species was randomly sampled. MRM tests assessed the beta diversity variance explained by host diet, geography, habitat, phylogeny, and “technical” parameters (see Supplemental Methods), with 4 beta diversity measures assessed: A) weighted UniFrac, B) unweighted UniFrac, C) Bray-Curtis, and D) Jaccard. Asterisks denote significance (adj. P < 0.05 for >95% of dataset subsets; see Methods). Beta diversity calculated on ASVs aggregated at the genus level. Box centerlines, edges, whiskers, and points signify the median, interquartile range (IQR), 1.5 × IQR, and >1.5 × IQR, respectively.

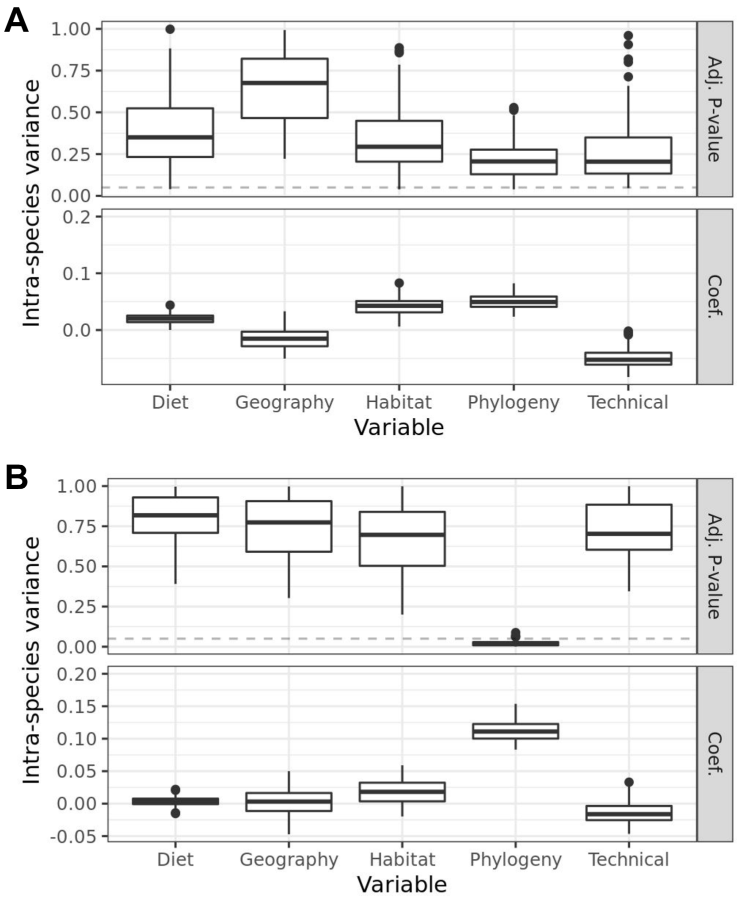


**Figure S10 .** *No host factors significantly explain archaeal alpha diversity*. The plots show the distribution of P-values (“Adj. P-value”) and partial regression coefficients (“Coef.”) across 100 dataset permutations used for multiple regression on matrix (MRM) tests. For each permutation, one individual per host species was randomly sampled. MRM tested whether inter-sample variance of alpha diversity was significant explained by host diet, geography, habitat, phylogeny, and “technical” parameters (see Methods), with 2 alpha diversity measures assessed: A) Shannon Index and B) Faith’s PD. No variables were significant (defined as adj. $P < 0.05$ for >95% of dataset permutations; see Supplemental Methods). Box centerlines, edges, whiskers, and points signify the median, interquartile range (IQR), 1.5 × IQR, and >1.5 × IQR, respectively.

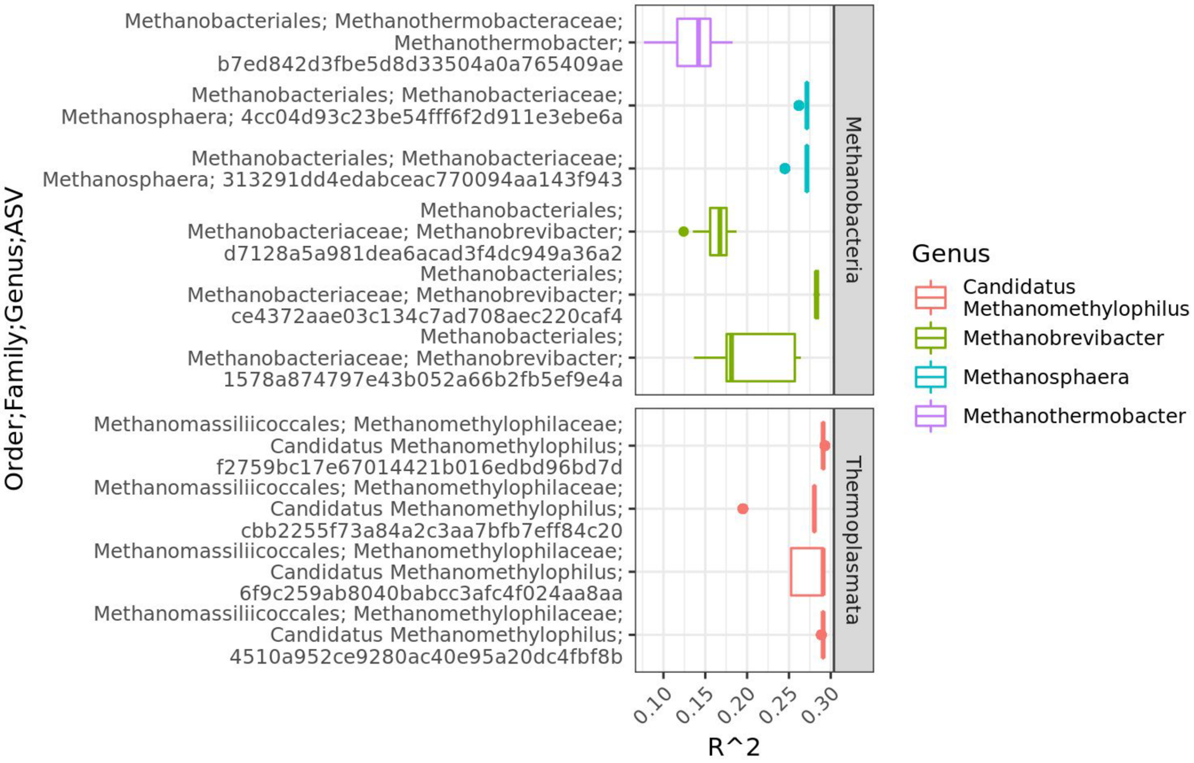


**Figure S11.** *Certain methanogen ASVs from multiple lineages are associated with diet, after accounting for host phylogeny*. Phylogenetic generalized least squares (PGLS) results for the ASVs with a significant association between ASV abundance and host diet, while accounting for host phylogenetic relatedness. Significance was defined as adj. $P < 0.05$ in ≥95% of permuted datasets, in which one sample per species was used per permutation. The boxplots depict the distribution of PGLS $R^2$ values across all 100 permutations. Box centerlines, edges, whiskers, and points signify the median, interquartile range (IQR), 1.5 × IQR, and >1.5 × IQR, respectively.

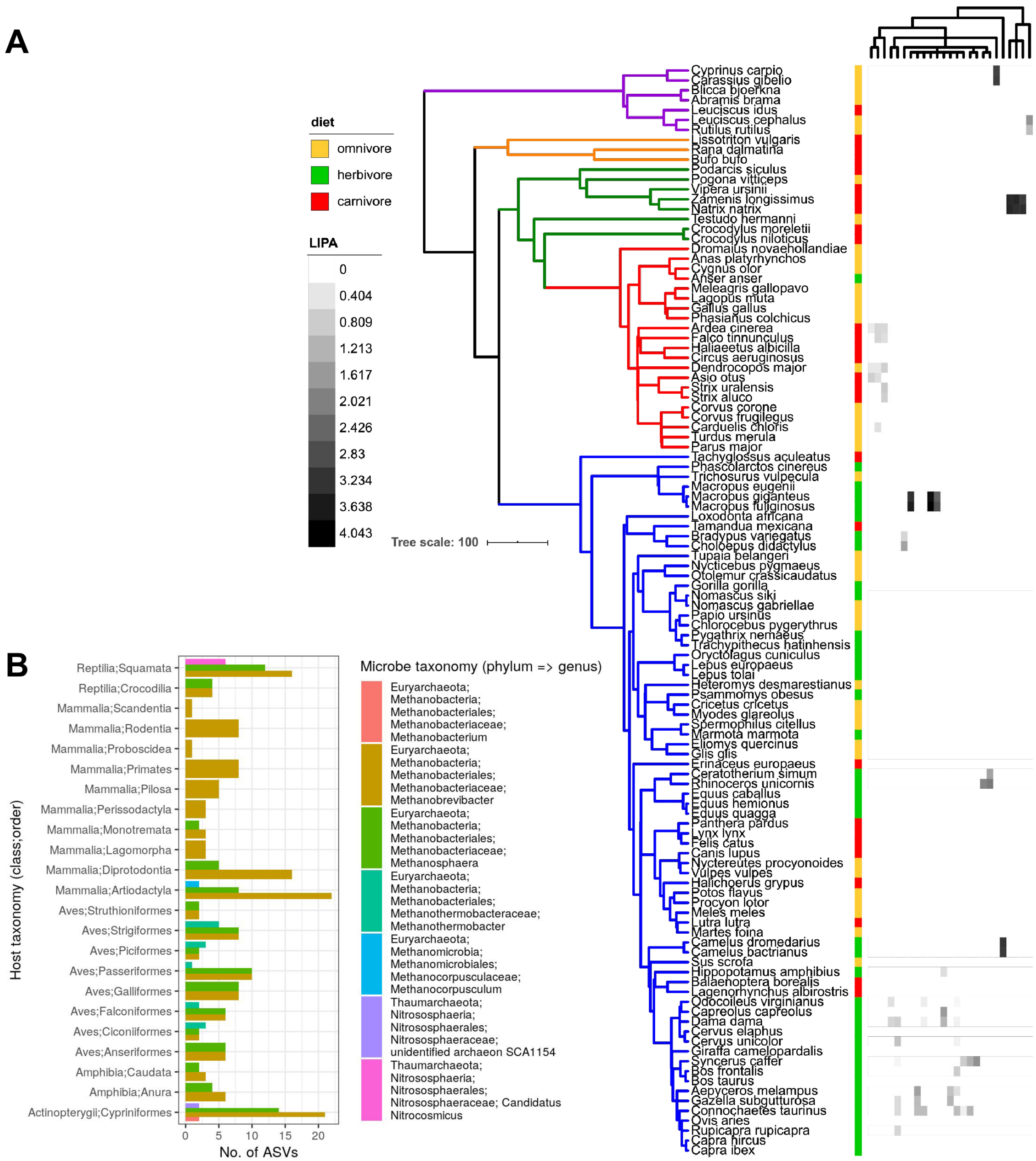


**Figure S12.** *The specific ASVs have similar abundances within certain vertebrate clades.* Various archaeal ASVs display local phylogenetic signal to various host clades. A) All ASVs with significant local phylogenetic signals (adj. $P < 0.05$) are mapped onto the host phylogeny. The phylogeny is the same as shown in Figure 1. The heatmap depicts local indicator of phylogenetic association (LIPA) values for each ASV–host association, with higher values indicating a stronger phylogenetic signal of ASV abundance. White boxes in the heatmap indicate non-significant LIPA tests. The dendrogram on the top of the heatmap is a cladogram based on taxonomy for each ASV (see Figure S13 for the full taxonomy). B) The bar plots show the number of ASVs with significant LIPA indices per archaeal genus and host clade.

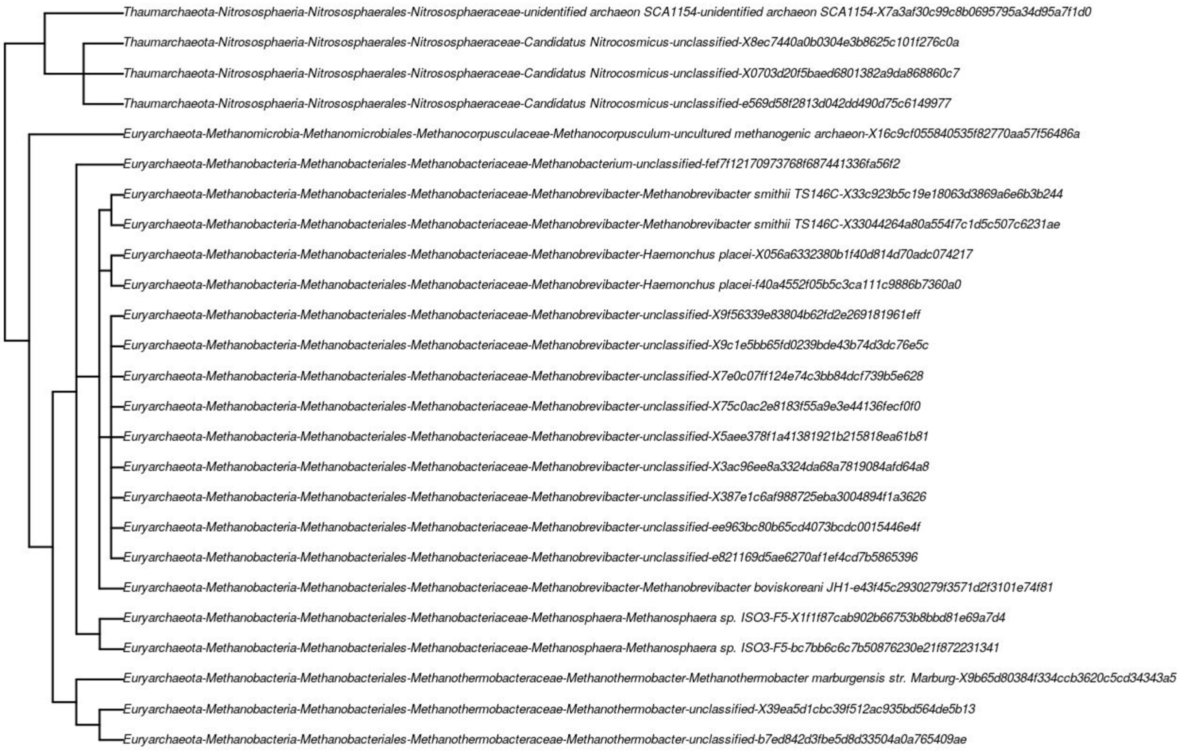


**Figure S13.** The cladogram as shown in Figure S12 with the entire ASV taxonomic classification as tip labels.

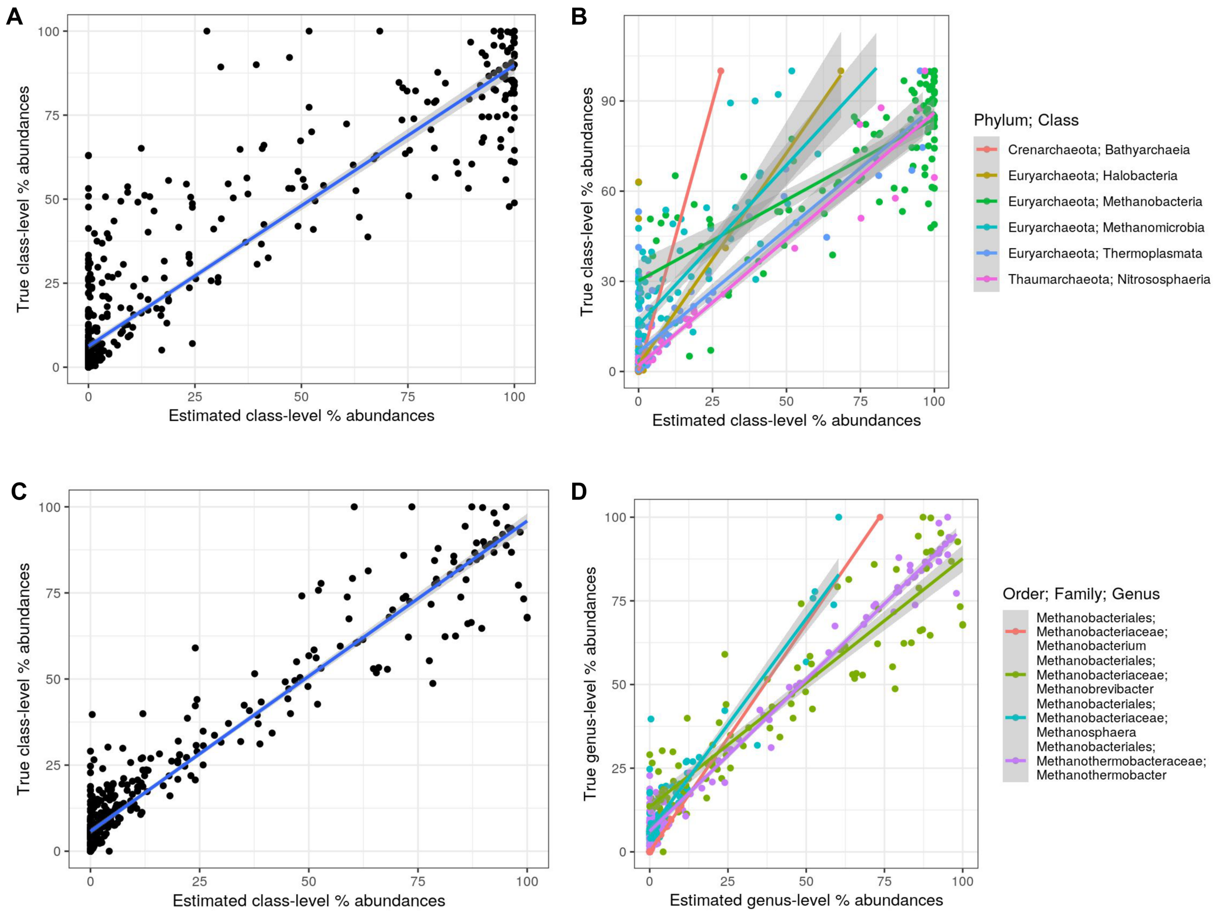


**Figure S14.** *Ancestral state reconstruction models accurately predict abundances in extant host species.* Linear regressions comparing ASR model predictions of archaeal abundances for each extant species relative to the observed mean abundance of all individuals per species. A) All class-level abundances, and B) abundances and linear regressions colored by class. C) All genus-level abundances for taxa belonging to Methanobacteria, and D) abundances and linear regressions colored by genus. Gray areas denote 95% confidence intervals for each linear model.

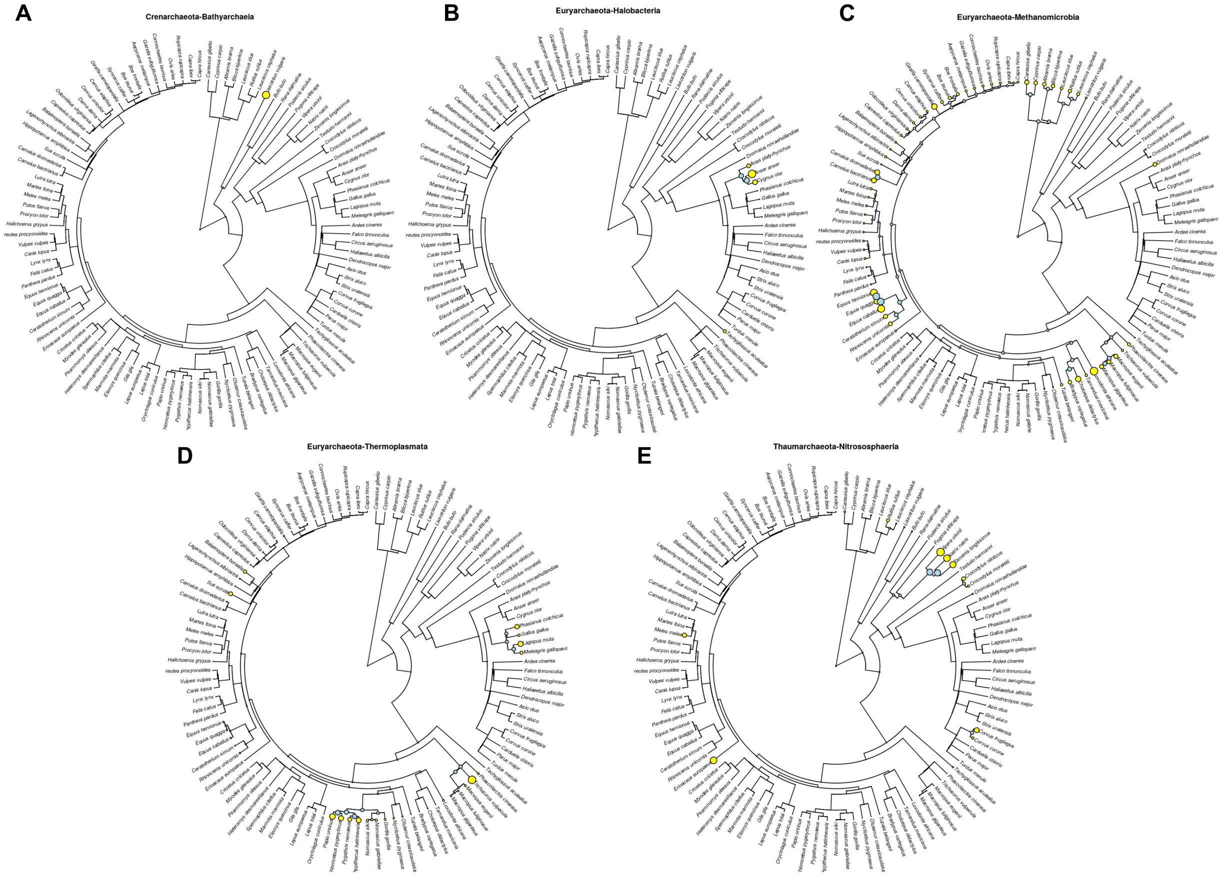


**Figure S15.** Predicted archaeal class-level abundance for extant host species (yellow circles) and and ancestral host species (blue circles): A) Bathyarchaeia, B) Halobacteria, C) Methanomicrobia, D) Thermoplasmata, and E) Nitrososphaeria. The phylogeny is the same as shown in Figure 1.

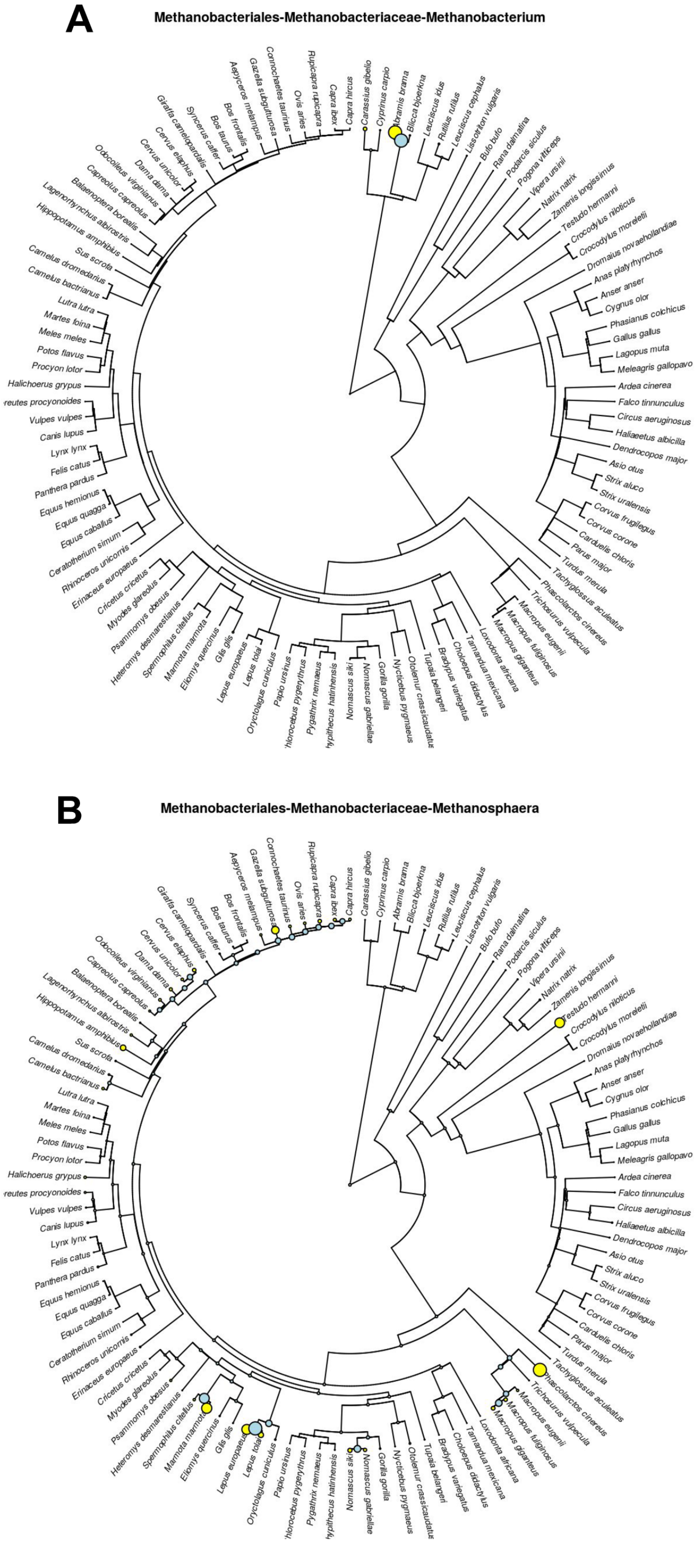


**Figure S16.** Predicted archaeal genus-level abundance for extant host species (yellow circles) and and ancestral host species (blue circles): A) Methanobacterium and B) Methanosphaera. The phylogeny is the same as shown in Figure 1.

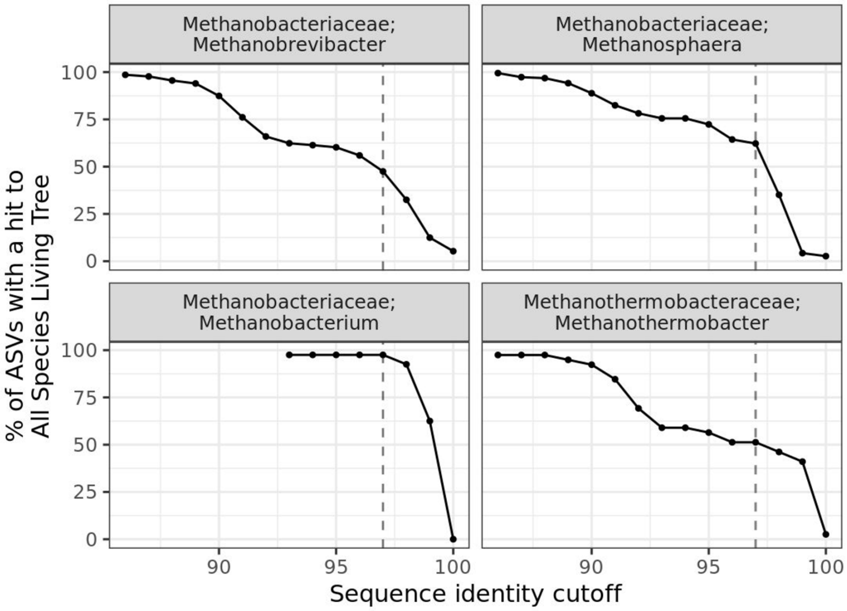


523 **Figure S17.** *Methanobacteria genera comprise a high proportion of uncultured ASVs.* Same as Figure S6, but just Methanobacteria genera. The plot facet labels are “family; genus”.

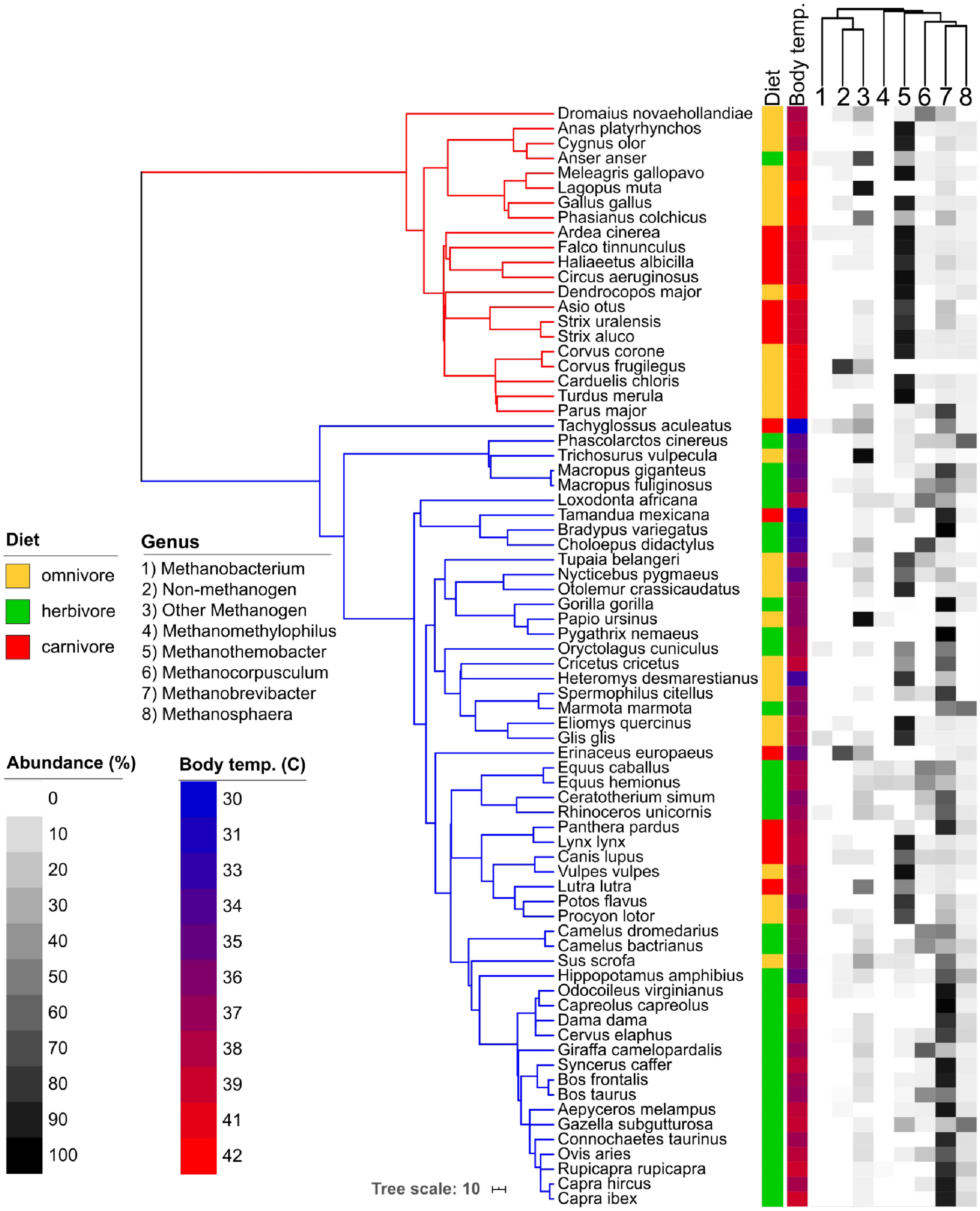


**Figure S18.** *Methanothermobacter is prevalent among avian species and associates with host body temperature.* The phylogeny is a pruned version (*n* = 74) of that shown in Figure 1. Host diet and body temperature are mapped into the tree along with genus-level archaeal abundances. The dendrogram above the heatmap is a cladogram depicting taxonomic relatedness. “Other Methanogen” refers to all other methanogen genera not specifically listed, and “Non-methanogen” refers to all non-methanogenic clades.

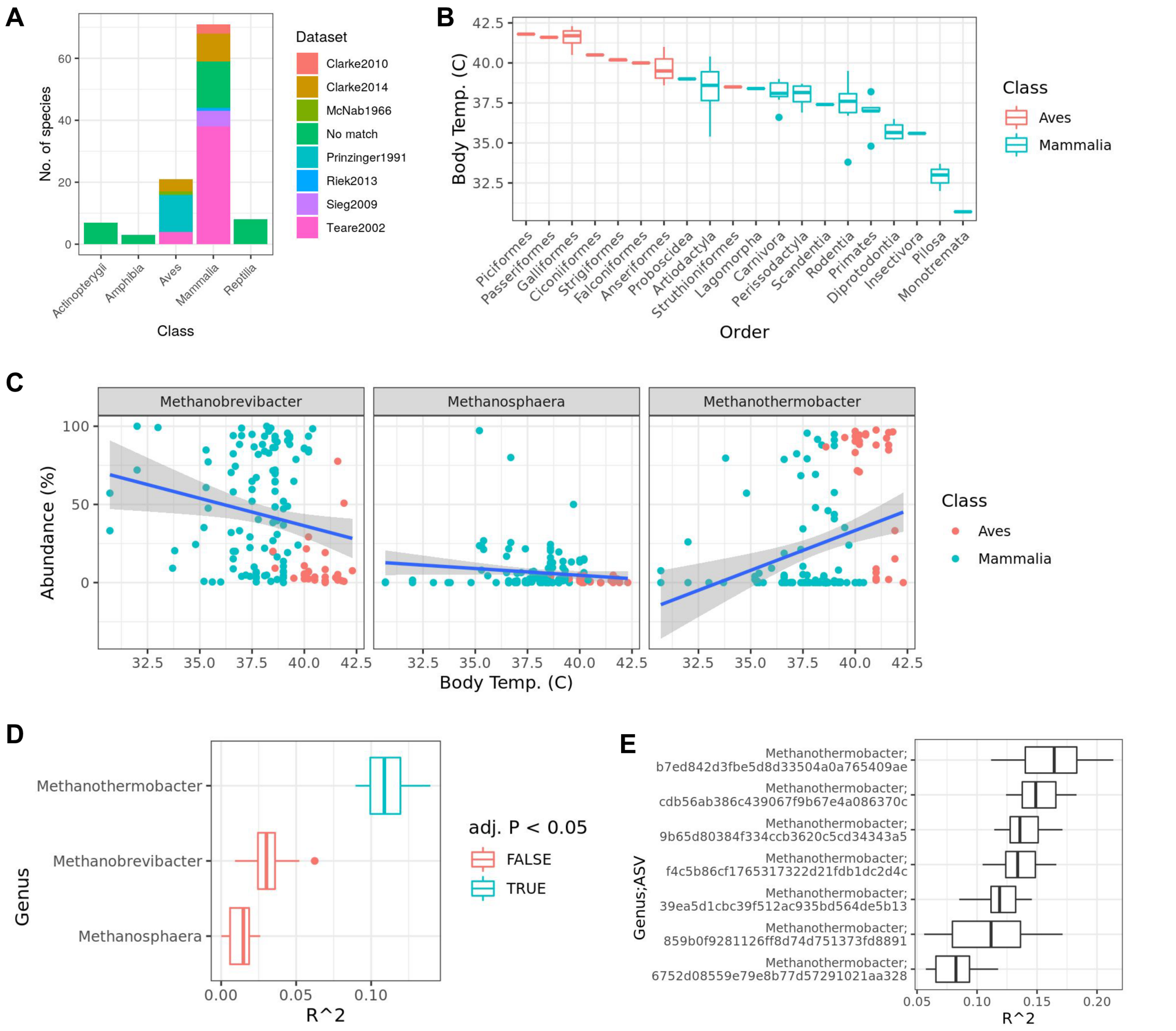


**Figure S19.** *Methanothermobacter abundance is explained by host body temperature*. A) The number of species with body temperature data, grouped by the body temperature dataset (see also Table S6). B) The distribution of body temperatures per host taxonomic order (one data point per species). C) Relative abundances of Methanobacteria genera as a function of host body temperature (celcius). The lines denote linear regressions with 95% CIs represented by the grey zones. D) RRPP coefficients of genus-level abundances as a linear function of host body temperature. Boxplots show the distribution across 100 permutations. E) The same as D, but ASV-level abundances used, with only significant ASVs shown. Note that host phylogeny was not used for the RRPP models shown in D & E. No taxa were significant when accounting for host phylogeny. Box centerlines, edges, whiskers, and points signify the median, interquartile range (IQR), 1.5 × IQR, and >1.5 × IQR, respectively.

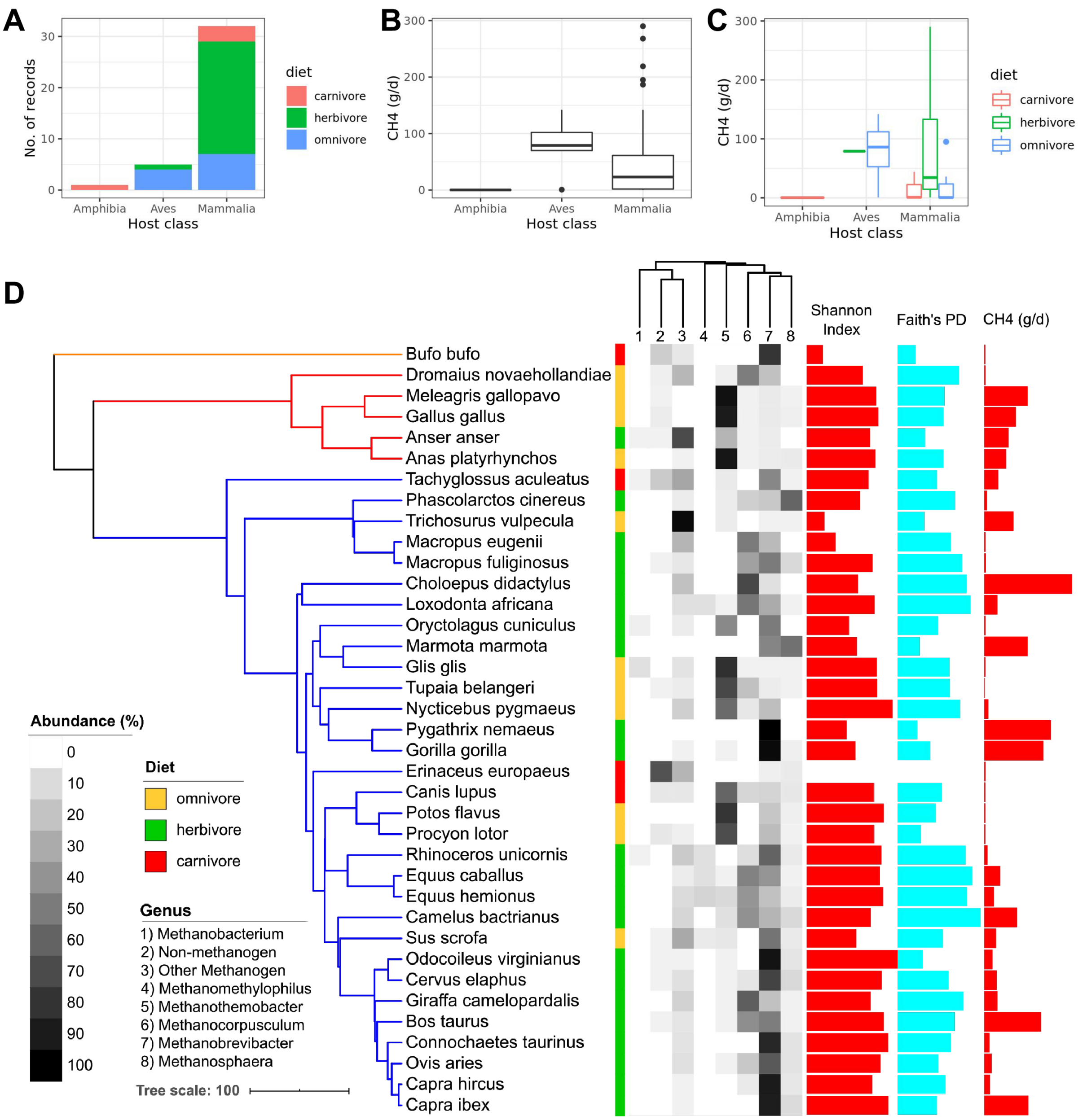


**Figure S20.** *Published animal methane emission data indicates that the avian species dominated by Methanothermobacter emit substantial amounts of methane.* A) The number of records obtained from Hackstein & van Alen 1996 ($n$ = 27) and Clauss et al., 2020 ($n$ = 10), grouped by host class and diet. B) & C) the distribution of methane emission rates per host species, grouped by class and C) colored by host diet. D) The phylogeny is a pruned version of that shown in Figure 1. From left to right, the data mapped onto the phylogeny is: host diet, methanogen genus mean abundances, methanogen ASV diversity (Shannon Index & Faith's PD), and methane emission rates. The lack of diversity values for *Erinaceus europaeus* (European hedgehog) is due to an absence of detectable methanogen ASVs. Box centerlines, edges, whiskers, and points signify the median, interquartile range (IQR), 1.5 × IQR, and >1.5 × IQR, respectively.

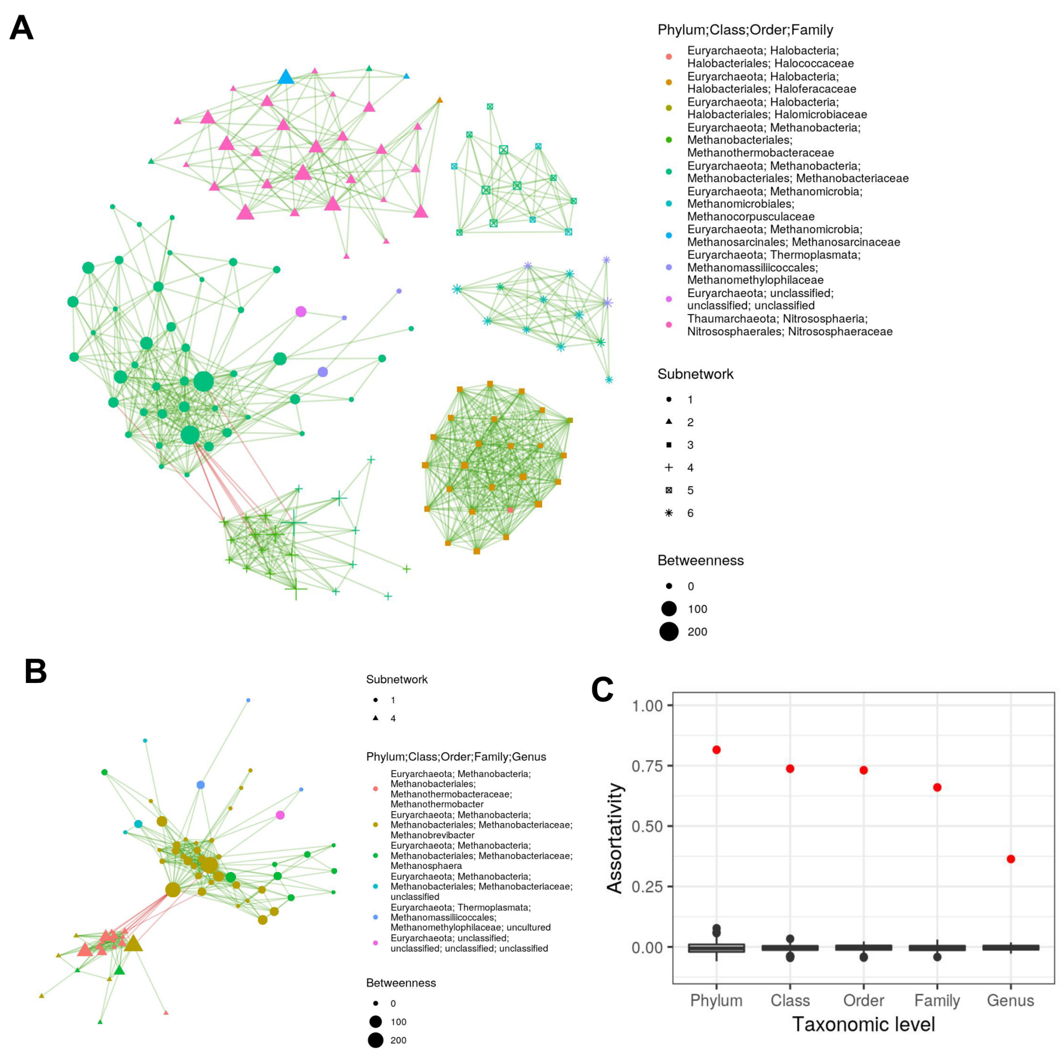


**Figure S21.** *Archaeal ASVs generally co-occur with members of the same taxonomic group.* The network nodes represent ASVs, with color denoting family-level taxonomic classifications, and shape denoting subnetwork (defined by clustering the network with the walktrap algorithm). Edges represent significant positive and negative co-occurrences among ASVs as denoted by green and red edges, respectively. Node size represents "betweenness", which is a measure of node connectedness. For clarity, only the largest 6 subnetworks are shown (but see Figure S22). B) Only subnetworks 1 and 4 are shown with node colors denoting genus-level classifications. C) The assortativity of ASVs by taxonomic level, in which a value of 1 means that all connected ASVs belong to the same taxonomic group, while a value of 0 denotes random association, and negative values indicate a dominance of inter-clade associations. The red points are the observed values, while the boxplots denote values for 100 permutations of networks with the same number of nodes and edges as the true network, but edges were randomly assigned. Box centerlines, edges, whiskers, and points signify the median, interquartile range (IQR), 1.5 × IQR, and >1.5 × IQR, respectively.

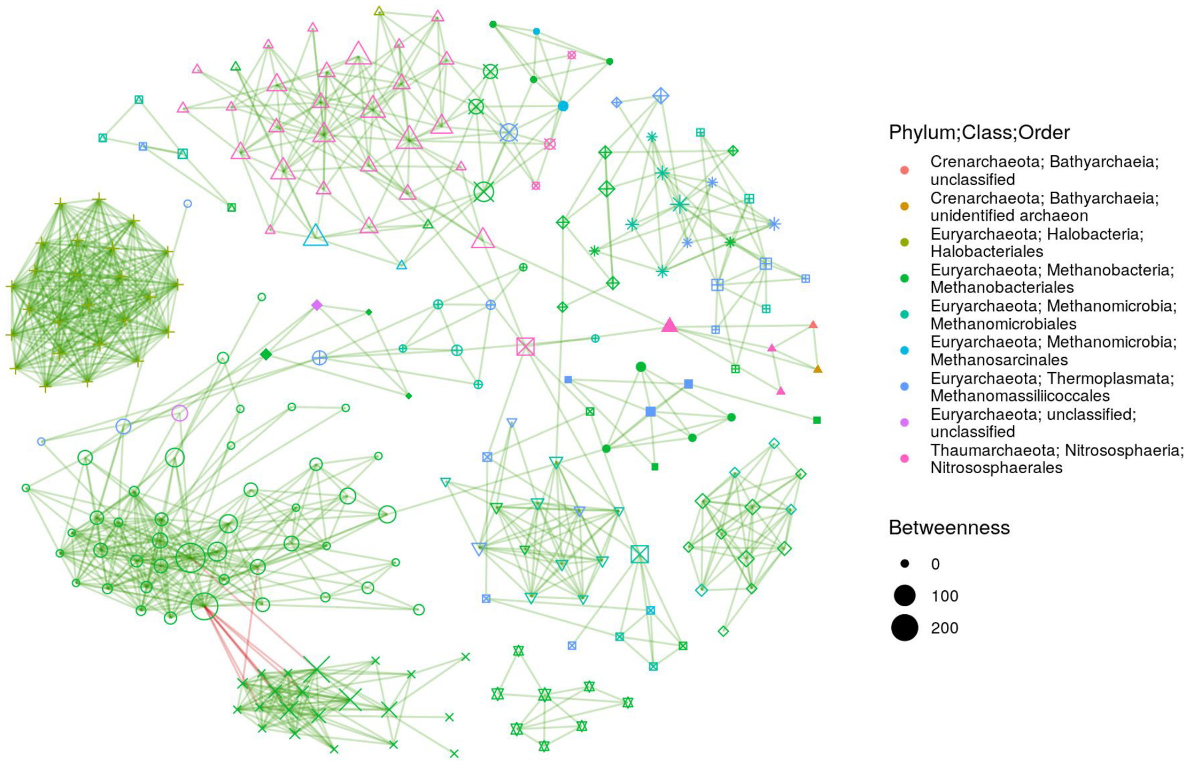


**Figure S22.** The same co-occurrence network as shown in Figure S21, but the largest 19 subnetworks are shown (238 of 313 ASVs) instead of just the largest 6 (151 of 313 ASVs). The entire co-occurrence network comprised 96 subnetworks, but to be able to distinguish among shapes denoting network nodes, only the top 19 subnetworks are shown.

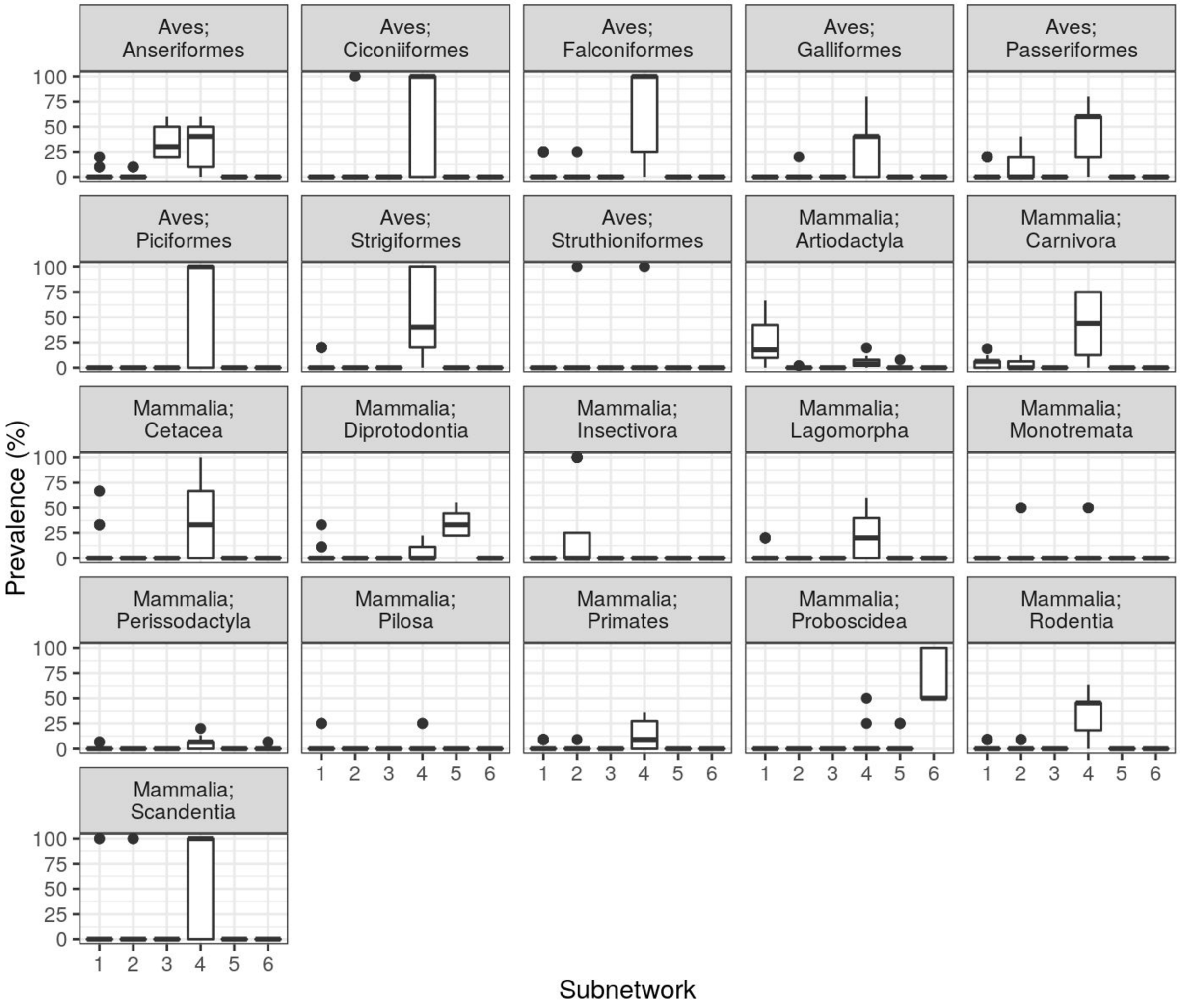


**Figure S23.** The percent of samples in which each ASV was observed (prevalence), grouped by the subnetwork to which each ASV belongs (see Figure S21A) and faceted by host taxonomic order. Box centerlines, edges, whiskers, and points signify the median, interquartile range (IQR), 1.5 × IQR, and >1.5 × IQR, respectively.

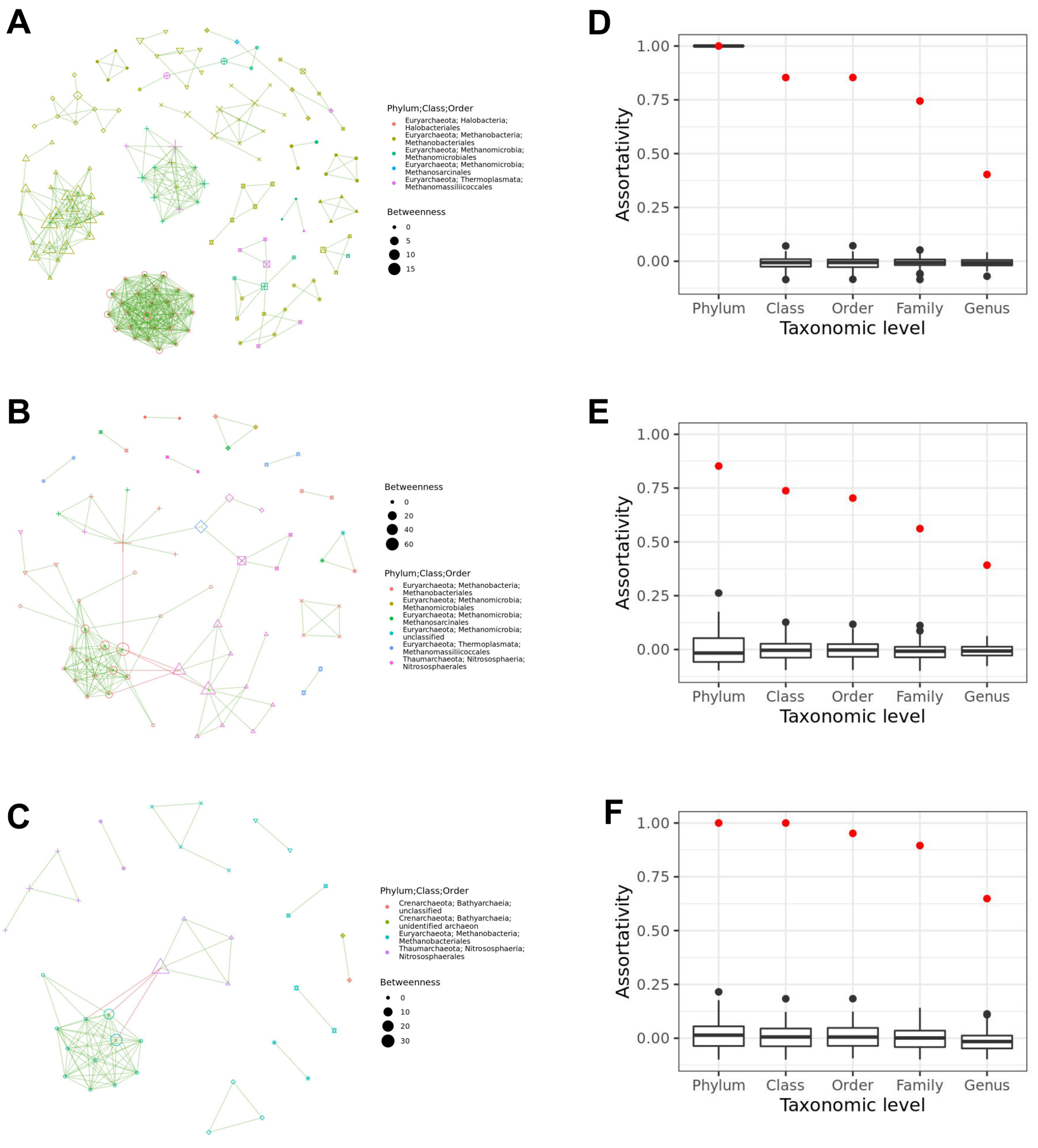


**Figure S24.** Co-occurrence networks for A) just herbivore, B) just omnivore, C) just carnivore samples. Node size represents “betweenness”, which is a measure of node connectedness. Green and red edges denote significant positive and negative co-occurrences, respectively. D-F) Assortativity of nodes the graph, determined for each taxonomic level from phylum to genus. High assortativity values indicate that the co-occurring taxa largely belong to the same taxonomic group. Box centerlines, edges, whiskers, and points signify the median, interquartile range (IQR), 1.5 × IQR, and >1.5 × IQR, respectively.

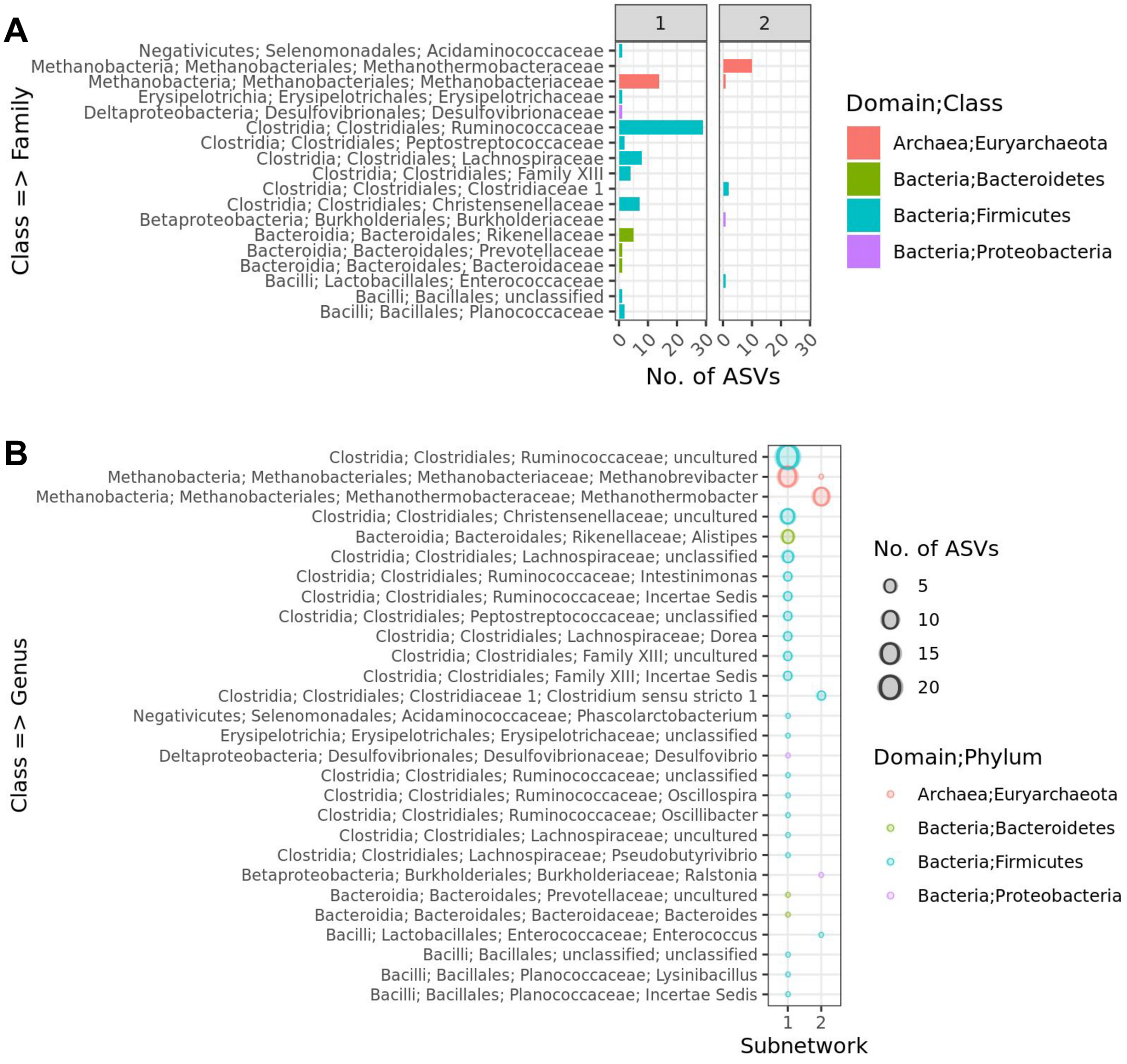


**Figure S25.** Taxonomic composition of the 2 sub-networks (Figure 4D) containing archaeal ASVs, with the number of ASVs summarized at the A) family and B) genus taxonomic levels.

Pausan, Manuela R., Cintia Csorba, Georg Singer, Holger Till, Veronika Schöpf, Elisabeth Santigli, Barbara Klug, Christoph Högenauer, Marcus Blohs, and Christine Moissl-Eichinger. 2019. “Exploring the Archaeome: Detection of Archaeal Signatures in the Human Body.” *Frontiers in Microbiology* 10 (December): 2796.

Pedersen, Thomas Lin. 2018a. “Ggraph: An Implementation of Grammar of Graphics for Graphs and Networks.” https://CRAN.R-project.org/package=ggraph. 2018b. “Tidygraph: A Tidy API for Graph Manipulation.” https://CRAN.R-project.org/package=tidygraph.

Pons, Pascal, and Matthieu Latapy. 2005. “Computing Communities in Large Networks Using Random Walks.” In *Computer and Information Sciences - ISCIS 2005*, 284–93. Springer Berlin Heidelberg.

Price, Morgan N., Paramvir S. Dehal, and Adam P. Arkin. 2010. “FastTree 2--Approximately Maximum-Likelihood Trees for Large Alignments.” *PloS One* 5 (3): e9490.

Prinzinger, R., A. Preßmar, and E. Schleucher. 1991. “Body Temperature in Birds.” *Comparative Biochemistry and Physiology. Part A, Physiology* 99 (4): 499–506.

Pruesse, Elmar, Christian Quast, Katrin Knittel, Bernhard M. Fuchs, Wolfgang Ludwig, Jörg Peplies, and Frank Oliver Glöckner. 2007. “SILVA: A Comprehensive Online Resource for Quality Checked and Aligned Ribosomal RNA Sequence Data Compatible with ARB.” *Nucleic Acids Research* 35 (21): 7188–96.

Quast, Christian, Elmar Pruesse, Pelin Yilmaz, Jan Gerken, Timmy Schweer, Pablo Yarza, Jörg Peplies, and Frank Oliver Glöckner. 2013. “The SILVA Ribosomal RNA Gene Database Project: Improved Data Processing and Web-Based Tools.” *Nucleic Acids Research* 41 (Database issue): D590–96.

Raymann, Kasie, Andrew H. Moeller, Andrew L. Goodman, and Howard Ochman. 2017. “Unexplored Archaeal Diversity in the Great Ape Gut Microbiome.” *mSphere* 2 (1). https://doi.org/10.1128/mSphere.00026-17.

R Core Team. 2020. *R: A Language and Environment for Statistical Computing*. Vienna, Austria: R Foundation for Statistical Computing.

Revell, Liam J. 2010. “Phylogenetic Signal and Linear Regression on Species Data: Phylogenetic Regression.” *Methods in Ecology and Evolution / British Ecological Society* 1 (4): 319–29.

Riek, Alexander, and Fritz Geiser. 2013. “Allometry of Thermal Variables in Mammals: Consequences of Body Size and Phylogeny.” *Biological Reviews of the Cambridge Philosophical Society* 88 (3): 564–72.

Schubert, Michael. 2019. “Clustermq Enables Efficient Parallelization of Genomic Analyses.” *Bioinformatics* 35 (21): 4493–95.

Sieg, Annette E., Michael P. O’Connor, James N. McNair, Bruce W. Grant, Salvatore J. Agosta, and Arthur E. Dunham. 2009. “Mammalian Metabolic Allometry: Do Intraspecific Variation, Phylogeny, and Regression Models Matter?” *The American Naturalist* 174 (5): 720–33.

Söllinger, Andrea, and Tim Urich. 2019. “Methylotrophic Methanogens Everywhere - Physiology and Ecology of Novel Players in Global Methane Cycling.” *Biochemical Society Transactions*, December. https://doi.org/10.1042/BST20180565.

Takai, K., and K. Horikoshi. 2000. “Rapid Detection and Quantification of Members of the Archaeal Community by Quantitative PCR Using Fluorogenic Probes.” *Applied and Environmental Microbiology* 66 (11): 5066–72.

Teare, Andrew. 2002. “International Species Information System, Medical Animal Records Keeping System (MedARKS) 2002 Data Extraction.” https://www.species360.org/about-us/mission-history/.

Thompson, Luke R., Jon G. Sanders, Daniel McDonald, Amnon Amir, Joshua Ladau, Kenneth J. Locey, Robert J. Prill, et al. 2017. “A Communal Catalogue Reveals Earth’s Multiscale Microbial Diversity.” *Nature* 551 (7681): 457–63.

Walters, William, Embriette R. Hyde, Donna Berg-Lyons, Gail Ackermann, Greg Humphrey, Alma Parada, Jack A. Gilbert, et al. 2016. “Improved Bacterial 16S rRNA Gene (V4 and V4-5) and Fungal Internal Transcribed Spacer Marker Gene Primers for Microbial Community Surveys.” *mSystems* 1 (1). https://doi.org/10.1128/mSystems.00009-15.

Wickham, Hadley. 2009. *ggplot2: Elegant Graphics for Data Analysis*. 1st ed. 2009. Corr. 3rd printing 2010 edition. New York: Springer.

Youngblut, Nicholas D., Georg H. Reischer, William Walters, Nathalie Schuster, Chris Walzer, Gabrielle Stalder, Ruth E. Ley, and Andreas H. Farnleitner. 2019. “Host Diet and Evolutionary History Explain Different Aspects of Gut Microbiome Diversity among Vertebrate Clades.” *Nature Communications* 10 (1): 2200.
